# CoRAL accurately resolves extrachromosomal DNA genome structures with long-read sequencing

**DOI:** 10.1101/2024.02.15.580594

**Authors:** Kaiyuan Zhu, Matthew G. Jones, Jens Luebeck, Xinxin Bu, Hyerim Yi, King L. Hung, Ivy Tsz-Lo Wong, Shu Zhang, Paul S. Mischel, Howard Y. Chang, Vineet Bafna

## Abstract

Extrachromosomal DNA (ecDNA) is a central mechanism for focal oncogene amplification in cancer, occurring in approximately 15% of early stage cancers and 30% of late-stage cancers. EcDNAs drive tumor formation, evolution, and drug resistance by dynamically modulating oncogene copy-number and rewiring gene-regulatory networks. Elucidating the genomic architecture of ecDNA amplifications is critical for understanding tumor pathology and developing more effective therapies.

Paired-end short-read (Illumina) sequencing and mapping have been utilized to represent ecDNA amplifications using a breakpoint graph, where the inferred architecture of ecDNA is encoded as a cycle in the graph. Traversals of breakpoint graph have been used to successfully predict ecDNA presence in cancer samples. However, short-read technologies are intrinsically limited in the identification of breakpoints, phasing together of complex rearrangements and internal duplications, and deconvolution of cell-to-cell heterogeneity of ecDNA structures. Long-read technologies, such as from Oxford Nanopore Technologies, have the potential to improve inference as the longer reads are better at mapping structural variants and are more likely to span rearranged or duplicated regions.

Here, we propose CoRAL (<u>Co</u>mplete <u>R</u>econstruction of <u>A</u>mplifications with <u>L</u>ong reads), for reconstructing ecDNA architectures using long-read data. CoRAL reconstructs likely cyclic architectures using quadratic programming that simultaneously optimizes parsimony of reconstruction, explained copy number, and consistency of long-read mapping. CoRAL substantially improves reconstructions in extensive simulations and 9 datasets from previously-characterized cell-lines as compared to previous short-read-based tools. As long-read usage becomes wide-spread, we anticipate that CoRAL will be a valuable tool for profiling the landscape and evolution of focal amplifications in tumors.

## Introduction

Oncogene amplification is one of the most common events in tumorigenesis contributing to tumor initiation and progression (Beroukhim et al. 2010; Steele et al. 2022). Often, these amplifications are mediated by the formation of circular, megabase-scale extrachromosomal DNA (ecDNA) (Turner et al. 2017; Wu et al. 2019; Kim et al. 2020). Previous studies have underscored the importance of ecDNA in driving tumor formation (Luebeck et al. 2023), evolution (Lange et al. 2022), oncogene-mediated gene regulation (Hung et al. 2021; Zhu et al. 2021), and drug resistance (Nathanson et al. 2014; Lange et al. 2022). Thus, profiling the genetic and structural landscape of small, focal amplifications (typically < 10Mb), such as ecDNA, in tumors is critical for understanding the mechanisms of tumor progression and developing more effective therapies.

Owing to the large and complex genomes of ecDNA, it remains challenging to accurately infer the set of “amplicon” structures present in tumors (Deshpande et al. 2019; Luebeck et al. 2020; Chapman et al. 2021). Existing approaches rely on paired-end short-read (Illumina) sequencing to identify amplicons from copy number profiles and breakpoints that then can be represented with an edge-weighted *breakpoint graph*; ecDNAs can subsequently be extracted as cycles from the breakpoint graph (Bafna and Pevzner 1996; Alekseyev and Pevzner 2009; Lin et al. 2014; Deshpande et al. 2019; Hadi et al. 2020). Despite the success of these approaches in predicting ecDNA presence in cancer samples (Deshpande et al. 2019; Kim et al. 2020; Luebeck et al. 2023), short-read reconstructions have several limitations. First, short-read approaches struggle to handle the highly-rearranged nature of ecDNA and accurately detect breakpoints, especially in repetitive or low-complexity regions. Second, because ecDNA can contain multiple copies of large segments that are unique in the reference (e.g., 1a), short-read data is limited in its ability to phase distant breakpoints correctly. Therefore, multiple collections of paths or cycles in the breakpoint graph can explain the increased copy-number equally well, masking the true structure (1c). Third, heterogeneity of ecDNA structures might result in multiple overlapping focal amplifications derived from the same genomic regions. To address these shortcomings of short-read technology, existing methods (e.g. AmpliconArchitect (AA) (Deshpande et al. 2019; Hung et al. 2022)) must use heuristics: for example, extracting cycles with the highest copy number iteratively from a breakpoint graph, until a large fraction of the aggregate copy number is explained. While these heuristic strategies return multiple small cycles (1c) that can later be recombined (Hung et al. 2022), they are still constrained by the intrinsic limitations of short-read technologies to identify structural variation and phase together distant breakpoints.

Long-reads have the potential to resolve these challenges. Recent research efforts utilized Oxford Nanopore reads to reconstruct simple ecDNA, building on off-the shelf *de novo* assembly (Helmsauer et al. 2020). However, *de novo* assembly methods often make choices based on underlying assumptions that do not hold: for example, they assume a diploid genome and that regions of high multiplicity are small enough to be spanned by long-reads. However, the heterogeneity of ecDNA structures violates the assumption of ploidy and the long segments of high multiplicity (typically 10kb-1Mb) in ecDNA are infrequently spanned by a single read, unlike the repetitive regions encountered in genome assembly, such as long interspersed nucleotide elements (LINEs) that are in the 10kb range. Concurrent with our method proposed below, a new approach, Decoil (Giurgiu et al. 2023), also aims at reconstructing ecDNA structures with long reads. However, it does not separate multiple distinct focal amplifications in one tumor sample, and uses a similar “simple cycle extraction and combining” heuristic designed for short read to reconstruct ecDNAs with high multiplicy segments. An alternative methodology utilizes optical mapping (Cao et al. 2014) (OM) to sequence large (> 200kbp) DNA fragments that span a limited number of the high multiplicity regions (Luebeck et al. 2020). While good for scaffolding, these data cannot precisely detect breakpoints, identify small structural variations, or resolve non-templated sequence, and work best in conjunction with short-read methods.

Here, we propose CoRAL (<u>Co</u>mplete <u>R</u>econstruction of <u>A</u>mplifications with <u>L</u>ong reads), an algorithm for reconstructing ecDNA amplicon sequence and structure from long-reads (such as those from Oxford Nanopore Technologies or PacBio). CoRAL builds a distinct breakpoint graph for each focally amplified region, and extract cycles (and walks) from the breakpoint graph representing ecDNA (and the potential focally amplified genomes). In cases where the reads are not always long enough to span the high multiplicity regions, CoRAL reconstructs likely cyclic architectures using quadratic programming that simultaneously optimizes parsimony of reconstruction, explained copy number, and consistency of long-read mapping. Through extensive benchmarks on simulated data and previously-characterized cell lines, we report that CoRAL substantially improves breakpoint detection and inferring the order of complex segments on ecDNA over long-read-based Decoil (Giurgiu et al. 2023) and the short-read-based AmpliconArchitect (Deshpande et al. 2019) methods.

**Figure 1:**
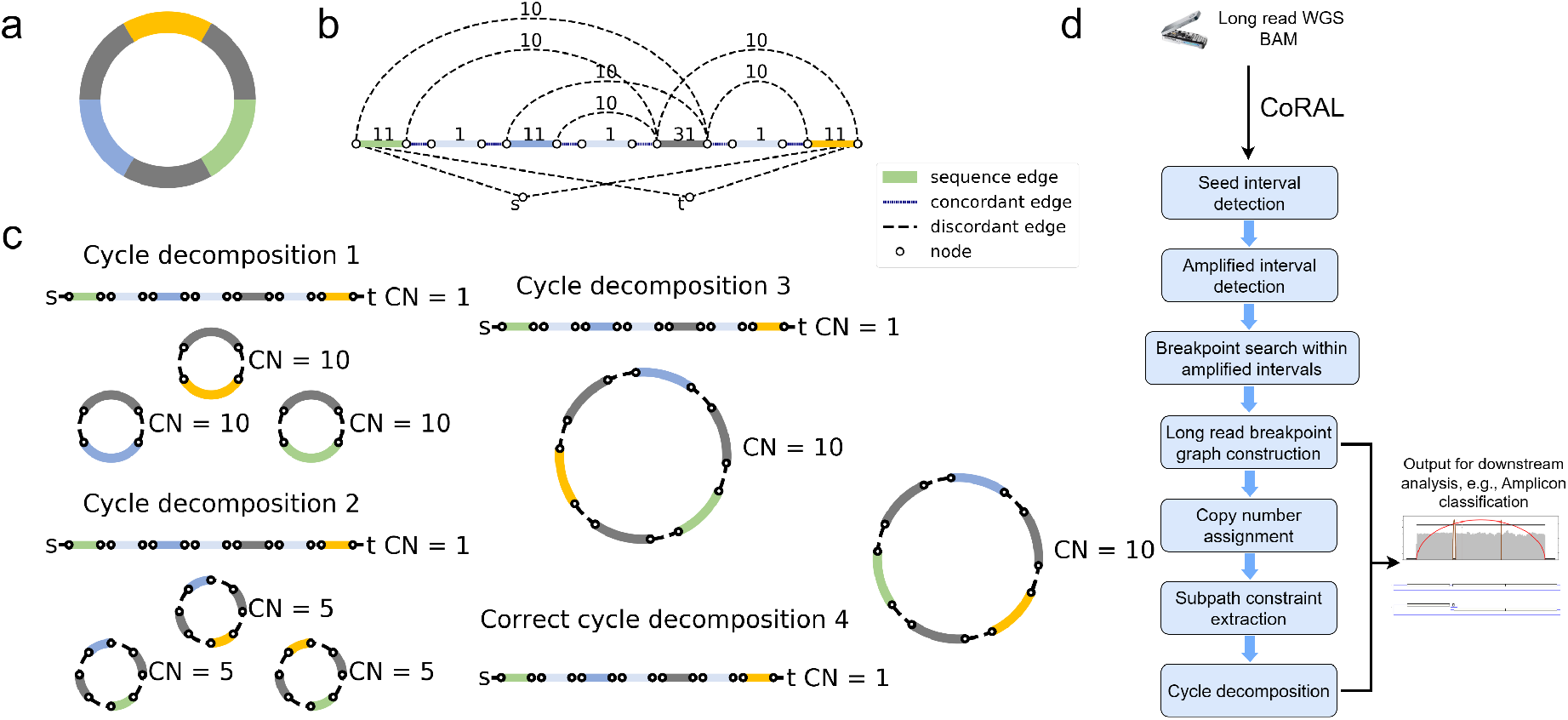
Long read based ecDNA reconstruction. (a) Native ecDNA structure, and copy number. (b) Cartoon of the breakpoint graph derived from the ecDNA in (a). Sequence edges represent segments of the reference genome. Concordant edges connect consecutive sequences with respect to the reference genome order, and discordant edges connect non-consecutive genome segments. Nodes are created at the endpoints of each sequence edge, and include source and sink nodes, s and t. (c) Multiple collections of decomposed paths and cycles from the breakpoint graph explain the changes in copy number and observed SVs. Long-reads that span regions of high multiplicity can help resolve the correct cycle. (d) Overview of the CoRAL method.

## Results

### An overview of the CoRAL method

For better exposition of the results, we first provide a brief description of the method. Details can be found in Methods and Appendix A2-A4. CoRAL takes mapped long-reads (in BAM format) as input and begins by identifying focally amplified *seed intervals*. The seed intervals can be provided directly, or derived from whole genome CNV calls (e.g., with third party tools like CNVkit (Talevich et al. 2016)) of mapped long reads. From the CNV calls, CoRAL selects genomic segments with minimum thresholds on copy number and aggregate size as seed intervals (Appendix A2).

CoRAL uses these seed intervals to construct a copy-number-weighted breakpoint graph separately for each amplified region. The graph construction starts with exploring all *amplified intervals* connected to the seed intervals through discordant edges given by chimeric long read mappings. Once all amplified intervals are identified for each focal amplification, a graph structure is organized by CoRAL to include the genome segments (sequence edges) from the amplified intervals, the concordant edges that join neighboring genome segments, and also the discordant edges within the amplified intervals and those connecting different amplified intervals. Once the graph structure is fixed, CoRAL recomputes the *copy number* for each edge which can best explain the long read coverage on each edge, while maintaining a balance of copy number between concordant and discordant edges incident on nodes. (see Methods and Appendix A3).

As its key step, CoRAL reconstructs potential ecDNA structures in the breakpoint graph by extracting a minimum number of *cycles and walks* from the graph, allowing duplication of nodes (e.g. 1c), where cycles represent the potential ecDNA species and walks represent linearly amplified or rearranged genome. Each cycle/walk is associated with a positive weight – corresponding to the copy number – so that the sum of length-weighted edges of extracted walks explains a large fraction of the total copy number of the edges in the breakpoint graph. In addition, CoRAL takes advantage of the fact that long-reads may span several breakpoints and incorporates these reads as *subwalk constraints*. In its cycle extraction, CoRAL also requires a majority of the subwalk constraints to be satisfied by the resulting cycles and walks, thus leveraging the power of long reads. CoRAL uses quadratic integer programming to solve a multi-objective optimization that minimizes the number of cycles/walks while maximizing the explained length-weighted copy number and the number of subwalk constraints (Methods and Appendix A4). It finally outputs the reconstructed breakpoint graphs for each focal amplification in the sample, as well as the associated cycles/walks from the graph. It also optionally outputs stylistic visualizations of the breakpoint graphs and cycles, as shown in subsequent results.

### Simulation benchmarks

We first assessed the effectiveness of amplicon reconstruction algorithms using simulated sequencing data from synthetic amplicon structures (Appendix A5, Supplementary Table 1, 2). To capture the diversity of ecDNA amplicons observed in patient tumors and cell lines, we simulated 75 distinct cyclic structures with varying numbers of breakpoints (between 1 and 20) from one of three origins: *episomal*, in which a contiguous region of the genome is excised from a chromosome; *chromothripsis*, in which a mitotic defect leads to the shattering of a lagging chromosome and ecDNA formation (Ly et al. 2017; Shoshani et al. 2021); or, finally, *2-foldback*, in which extruding double-stranded DNA from a stalled replication fork is broken off as ecDNA (Passananti et al. 1987). Our simulated ecDNAs additionally included internal structural variants in the form of insertions, deletions, duplications and inversions (see Appendix A5 for more detailed description of the simulation process and Supplementary Table 1 for the data). Subsequently, each test dataset was generated by randomly selecting between 1 and 5 amplicon structures (from the pool of 75 synthetic amplicons). Reads from long-read (using Nanosim (Yang et al. 2017)) and Illumina short-read, paired-end technologies (using Mason (Holtgrewe 2010)) were simulated from these amplicons at one of three coverages (50X, 100X, or 250X coverage; or approximate copy-numbers of 7, 15, or 37, respectively) and merged with reads from one of five simulated normal, diploid genomes (each with ∼13X coverage). A total of 50 test datasets were simulated in this fashion and used for benchmarking amplicon reconstruction (Supplementary Table 2).

From these inputs, ecDNA was reconstructed using simulated long-reads provided to CoRAL and Decoil (Giurgiu et al. 2023) - a separate long-read amplicon reconstruction tool - or simulated short-reads provided to AmpliconArchitect (AA). In most cases, the *heaviest* CoRAL cycle, defined as the cycle with the largest length-weighted copy number, was better at recapitulating the true architecture compared to the AA cycle (e.g., 2a). We systematically evaluated the accuracy of the best reconstruction *W*_*r*_ (as defined as the highest-scoring reconstruction with respect to a particular statistic) against a true cycle *W*_*t*_ using four additional measures defined briefly below (2b-e; see Appendix A7 for more detailed definitions):

**Figure 2:**
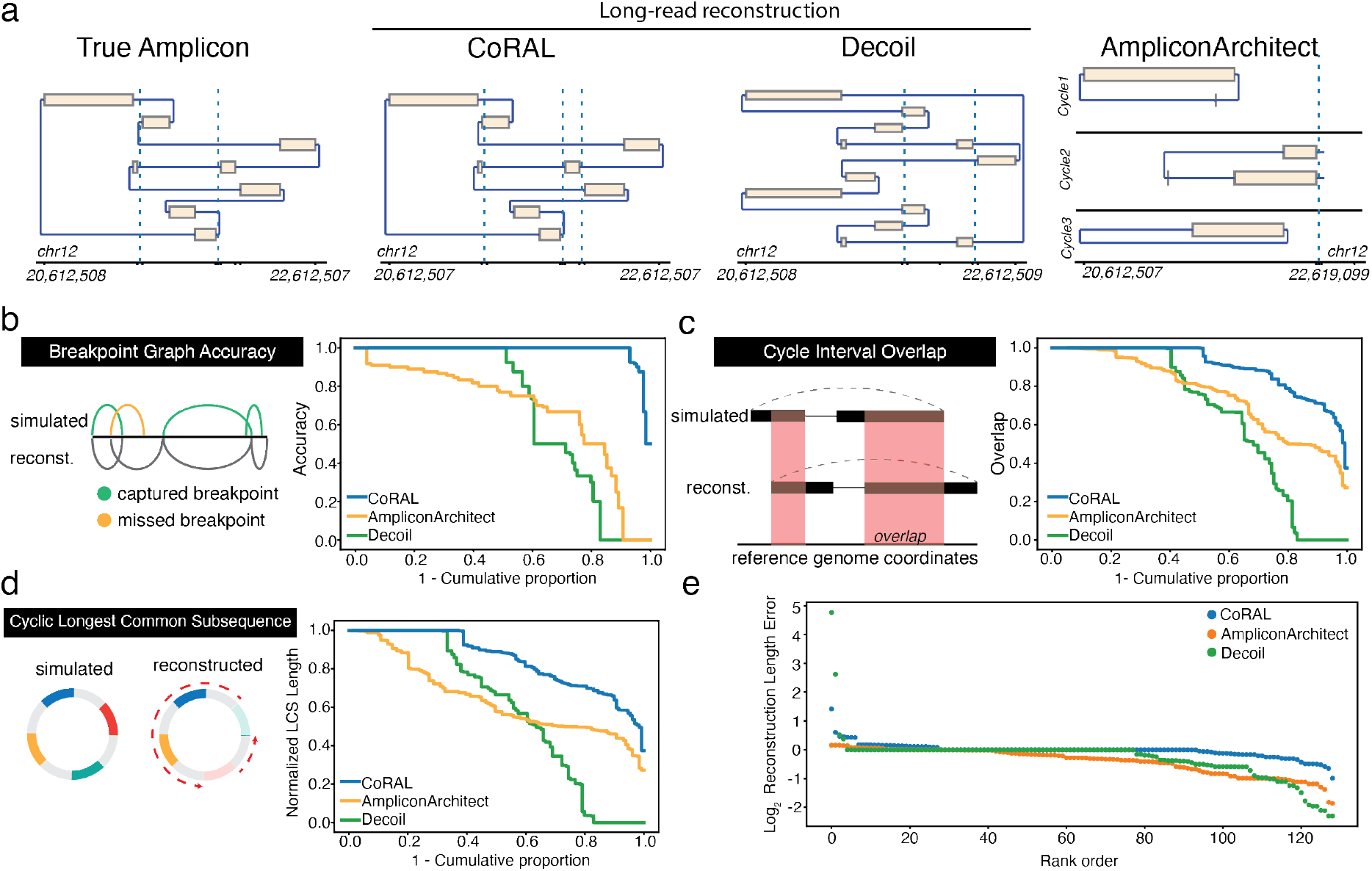
Overview of simulation benchmarking. (a) True structure compared to CoRAL, Decoil, and AA reconstructions for an example amplicon (Episomal, 8 observed breakpoints). (b-e) Cumulative distributions of CoRAL, Decoil, and AA reconstructions across all simulations for (b) breakpoint graph accuracy, (c) cycle interval overlap, (d) cyclic longest common subsequence and (e) rank-order distribution of reconstruction length error. Empirical cumulative densities are reported for (b), (c) and (d); and each point in (e) corresponds to a simulated amplicon. See Appendix A7 for more detailed information.

1. **Breakpoint Graph Accuracy** reports the proportion of discordant edges that agree, up to a tolerance of 100bp, between the true breakpoint graph *𝒢*_*t*_ and reconstructed breakpoint graph 𝒢_*r*_.
2. **Cycle Interval Overlap** measures the Jaccard index, weighted by the number of nucleotides, of the genomic intervals defined by *W*_*t*_ and *W*_*r*_.
3. **Cyclic Longest Common Subsequence (LCS)** measures the length of the longest common subsequence contained in *W*_*t*_ and *W*_*r*_ *after* eliminating intervals that are not found in both, normalized to the length of *W*_*t*_.
4. **Reconstruction length error** reports the difference in amplicon lengths between *W*_*r*_ and *W*_*t*_, normalized by the true amplicon length *W*_*t*_. We report *log*_2_-scaled values.

Across the 50 simulated datasets, we observed consistently improved reconstruction of CoRAL over AA and Decoil for all four measures (2b-e). Notably, 93% of CoRAL reconstructions perfectly recapitulated the ground truth breakpoint graph as compared to 51% for Decoil and 4% for AA (2b). These results underscore the improved mapping of structural variants with long-reads.

While CoRAL outperformed Decoil and AA in all four measures, both AA and Decoil capture many critical aspects of the amplicon, such as including the most amplified intervals (2c) and capturing the true ordering of the segments (2d). Mostly, CoRAL’s improved performance is reflected in reconstructed cycle lengths that are most similar to the true cycle (2e). In addition, we observe that both AA and CoRAL tend to produce a main cycle that account for a large fraction of length-weighted copy-number (Supplementary Fig. 1) and that this weight ratio is correlated with cycle reconstruction accuracy (Supplementary Fig. 2). Through these analysis, we also noted several examples where the interval ordering is incorrect despite near-perfect recovery of breakpoint graph and interval overlap (Supplementary Fig. 3), reflecting the technological limitations of reads that were not long enough to resolve the true order of segments.

We also compared reconstruction performance as a function of the complexity of amplicons (number of segments, or sequence edges), sequence coverage, their formation context, and level of duplication (or multiplicity). We observed that the number of segments in the true amplicon had modest effects on reconstruction accuracy, (Supplementary Fig. 4), as did coverage (and by extension copy-number; Supplementary Fig. 5). These observations suggest that all algorithms, but especially CoRAL, can accurately reconstruct complex ecDNAs at low copy numbers (e.g. < 7). We additionally observed that increasing levels of segmental or breakpoint multiplicity often resulted in poorer reconstruction accuracy for all methods tested, though CoRAL remained mostly robust (Supplementary Fig 6). However, in considering the various contexts in which ecDNA can form (Bafna and Mischel 2022), we observed substantial performance differences: generally, we observed that chromothripsis amplicons were most difficult for AA and CoRAL with Decoil modestly outperforming CoRAL conversely 2-foldback amplicons were most difficult for Decoil. These observations highlight the importance of accurately detecting structural variants, which is greatly enhanced with long reads, but can be nevertheless challenging depending on the complexity of breakpoints (Supplementary Fig. 7).

### Amplicon reconstruction in cell lines

Next, we evaluated amplicon reconstruction using matched Nanopore long-read sequencing and Illumina paired-end short-read sequencing in 7 previously characterized cell-lines spanning a range of cancer types and amplifications (summarized in Table 1 and Supplementary Table 3): COLO320(- DM, -HSR), PC3(-DM, -HSR), GBM39(-HSR), and CHP-212. Of these 7 cell lines, there are 3 isogenic pairs in which the amplified oncogene is located on chromosomal homogeneously staining regions (HSRs) while maintaining the core cyclic structure, as opposed to ecDNA (e.g., COLO320-HSR vs COLO320-DM). Additionally, we assessed reconstruction in 4 recently monoclonalized versions of these cell lines (PC3-DM, PC3-HSR, GBM39ec, and COLO320-DM). Together, this resulted in a matched Nanopore and Illumina data for 10 samples for analysis.

**Table 1:** Overview of cell lines profiled in this study. Cell type name, cancer type, subset of important amplified oncogenes, amplification type, monoclonal status, and source. Abbreviations: HSR = Homogeneously Staining Region; ecDNA = extrachromosomal DNA.

| Cell line | Cancer type | Gene(s) | Amp. | Mono. | Source |
| --- | --- | --- | --- | --- | --- |
| PC3DM (mono) | Prostate | *MYC* | ecDNA | Yes | This study |
| PC3HSR (mono) | Prostate | *MYC* | HSR | Yes | This study |
| GBM39 (mono) | Glioblastoma | *EGFRvIII* | ecDNA | Yes | This study |
| COLO320-DM (mono) | Colorectal | *MYC* | ecDNA | Yes | This study |
| COLO320-DM | Colorectal | *MYC* | ecDNA | No | Hung 2021<br>(Hung et al. 2021) |
| COLO320-DM | Colorectal | *MYC* | ecDNA | No | Wu 2019<br>(Wu et al. 2019) |
| GBM39 | Glioblastoma | *EGFRvIII* | ecDNA | No | Wu 2019<br>(Wu et al. 2019) |
| CHP-212 | Neuroblastoma | *MYCN, TRIB2* | ecDNA | No | Helmsauer 2020<br>(Helmsauer et al. 2020) |
| COLO320-HSR | Colorectal | *MYC* | HSR | No | Wu 2019<br>(Wu et al. 2019) |
| GBM39-HSR | Glioblastoma | *EGFRvIII* | HSR | No | Wu 2019<br>(Wu et al. 2019) |

### CoRAL accurately predicts the existence of ecDNA

We ran the AmpliconClassifier (Luebeck et al. 2023) method to reconfirm the cyclic structure of the ecDNA amplicons in all samples. AmpliconClassifier parses the breakpoint graph and identifies sub-graphs as being cyclic (or ecDNA), breakage fusion bridge, heavily-rearranged, or linear-rearranged (Kim et al. 2020). CoRAL identified altogether 60 amplicons in the 10 cell lines, including the main ecDNA (or HSR) amplicon in each sample. AmpliconClassifier consistently classified the main ecDNA amplicon as cyclic with the breakpoint graphs constructed by CoRAL using long-reads and AA using short-reads, indicating the existence of ecDNA (or HSR). Long-read sequencing did not identify new ecDNA amplicons nor fail to detect previously confirmed ecDNA amplicons in the cell line samples.

### CoRAL cycles better explain the copy numbers in ecDNA amplicons

To benchmark the reconstruction quality of CoRAL and AA in cell lines, we computed the fraction of length weighted copy numbers in the breakpoint graph given by the *k*-heaviest cycles, which we previously observed correlated with accuracy (with *k* = 1), for *k* = 3 and *k* = 1, in each of the 10 ecDNA cell lines (3a, Supplementary Fig. 8a; see Appendix A7 for details of the statistic). Consistent with simulated data, these results demonstrated that CoRAL explains a higher fraction of the length-weighted copy number with fewer cycles. Across all samples, the copy number explained by the 3 heaviest cycles was substantially higher for CoRAL compared to AA (3a). The reconstructed cycles of COLO320-DM (Wu2019), the monoclonal COLO320-DM (mono) and the shallow coverage COLO320-DM (Hung2021) showed consistent heaviest cycle weight ratio (3a, Supplementary Fig. 8a), shared many structural features, and contained a similar subset of genes (3c,d, Supplementary Table 4), even as they showed some differences in the reconstructed amplicons (Supplementary Figure 9). These differences could reflect differences in intrinsic heterogeneity or evolution of the cell line over time, which also resulted in lower heaviest cycle weight ratio in COLO320-DM cells and its isogeneic pair COLO320-HSR, in comparison to the GBM39 and CHP-212 cell lines with a single dominating ecDNA structure (Wu et al. 2019; Helmsauer et al. 2020) (Supplementary Fig. 8a).

### CoRAL cycles satisfy more subwalk constraints

In its optimization, CoRAL takes advantage of the fact that long-reads may span several breakpoints and incorporates these reads as *subwalk contraints* that can be satisfied during cycle decomposition and lend support for accurate reconstruction. As such we mapped each long read subwalk constraint to each AA cycle and checked if the subwalk constraint can also be satisfied by that cycle. Expectedly, CoRAL satisfied more subwalk constraints compared to AA (3b, Supplementary Fig. 8b), especially for the complex amplicons. For example, CoRAL satisfied 1.5x and 25x more subwalk constraints in COLO320-DM (mono) and PC3-DM (mono), respectively. Together, these subwalk constraints support most junctions of the amplicon (e.g., see 3c,d), thereby taking advantage of the long-reads that span multiple breakpoints. Nevertheless, no reconstruction satisfied all subwalk constraints in either CoRAL or AA, consistent with the high heterogeneity of ecDNA structure in samples.

**Figure 3:**
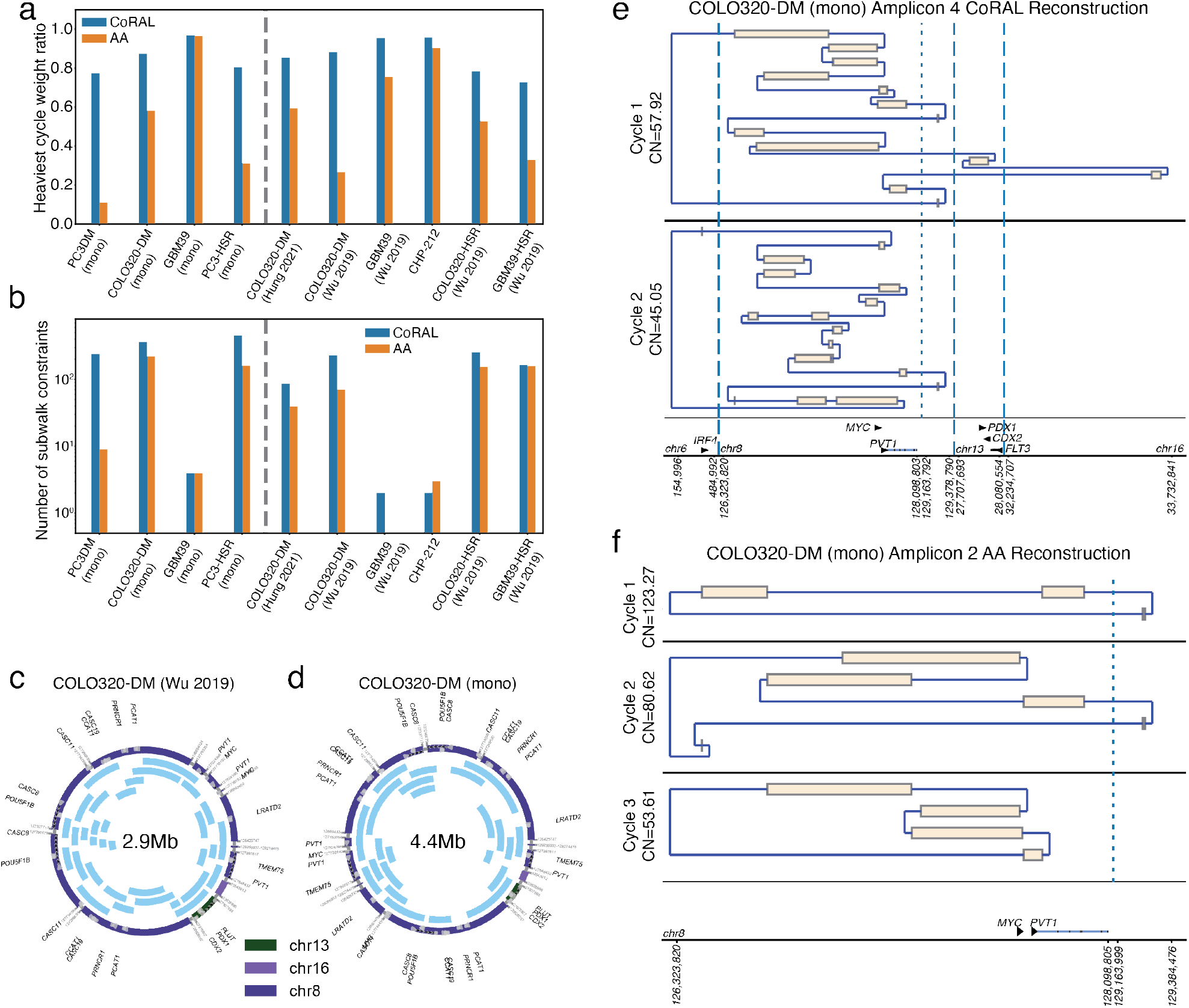
Amplicon reconstruction in cell lines. (a) Fraction of length-weighted copy numbers given by the 3 heaviest cycles, reported by CoRAL and AA; (b) number of satisfied subwalk constraints by CoRAL and AA, in cell lines. (c) The cycle with largest length-weighted CN from previously published COLO320-DM. (d) The cycle with largest length-weighted CN from monoclonalized COLO320-DM. In (c) and (d), each black arc within the cycle indicates a subwalk constraint satisfied by that cycle. (e) The two cycles with largest length weighted copy numbers by CoRAL. (f) The three cycles with largest length weighted copy numbers by AA.

### CoRAL cycles enable the study of critical aspects of amplicon structures

Reconstruction supported by long-read subwalk constraints additionally enabled the study of critical aspects of the amplicon structures. As one example, 3e,f shows the reconstruction of the two heaviest CoRAL and the three heaviest AA cycles, respectively, for monoclonal COLO320-DM. To note, the monoclonal COLO320-DM is a recently derived line from a parental line where previous experiments integrating WGS, optical mapping, and in-vitro ecDNA digestion revealed an ecDNA structure of approximately 4.3Mb (Hung et al. 2021). Here, the automated reconstruction of monoclonal COLO320-DM using CoRAL also revealed an ecDNA of size 4.4Mbp (3d) which shared many structural features with the previous reconstruction.

One distinct feature of the COLO320-DM *MYC* amplicon is the overexpression of a fusion transcript consisting of a truncated, 5’ portion of the lncRNA *PVT1* fused to the second exon of the *MYC* oncogene (Hung et al. 2021). This is despite *PVT1* being positioned downstream of *MYC* in the reference genome. As expected, CoRAL reconstruction of COLO320-DM includes a breakpoint that connects a truncated, 5’ portion of *PVT1* upstream of exon 2 of *MYC*, thereby explaining the fused transcript. Notably, both CoRAL and AA detected the *PVT1* -*MYC* fusion breakpoint in all COLO320-DM samples; however, CoRAL’s cycle decomposition included this breakpoint in the heaviest (largest length-weighted CN) cycle across multiple COLO320-DM samples (3c, d, e, Supplemetary Table 4). AA did not include the breakpoint in the 3 heaviest cycles (3f), instead, it reports a smaller cycle of size ∼ 90 kbp containing the breakpoint by itself. Furthermore, subwalk constraints due to long-reads linked truncated *PVT1* and *MYC* on a single molecule. Correspondingly, CoRAL reconstructions of cycles in COLO320-DM (mono), COLO320-DM (Hung 2021), and COLO320-DM (Wu 2019) all showed the three elements in a single cycle (3c, d, Supplementary Table 4).

We additionally observed that subwalk constraints and CoRAL’s cycle reconstructions support a co-amplification of the ncRNA *PCAT-1* and *MYC* on COLO320-DM ecDNA (3e,f). Previous DNA FISH experiments also confirmed the co-existence of these genes on COLO320-DM (Hung 2021 (Hung et al. 2021), Extended Figure 4g). The *PCAT-1* ncRNA is known to repress *BRCA2* (Prensner et al. 2014b), activate *MYC* (Prensner et al. 2014a), and promote cell proliferation (Xiong et al. 2019), and is upregulated in prostate, colorectal, and other cancers (Xiong et al. 2019). Thus, these CoRAL cycle reconstructions are also consistent with the regulatory and pro-oncogenic roles of *MYC* and *PCAT-1*. Together, these results highlight the advantages of CoRAL in reconstructing complex ecDNAs in cell lines and may enable new biological insights into the co-amplification of genetic elements on the same ecDNA molecule.

### CoRAL requires comparable computational resources to AA

We finally compared the computational resources required by CoRAL and AA to reconstruct all amplicons in these cell lines. To perform a fair test, we ran CoRAL and AA on the same Ubuntu system (2x Intel Xeon X5680 CPUs, and 128G RAM). Importantly, we observed that total running time and memory of CoRAL was comparable to AA for reconstructing the amplicons, even if an MIQCP was solved for each amplicon (Supplementary Figure 10). The most complex sample, COLO320-HSR (Wu 2019), was completed in less than 22h (∼ 8 * 10^5^s) for CoRAL. Furthermore, we found that most focal amplifications except ecDNA are relatively easy to resolve, with the resulting breakpoint graphs being small – out of the 60 amplicons detected by CoRAL across all samples, only 8 required greedy MIQCP, including 7 of the 10 total ecDNA amplicons.

## Discussion

Our results suggest that long-read guided reconstruction greatly improves ecDNA structure resolution, both in individual detection of breakpoints and in the accuracy of the large-scale predicted structure. The constrained optimization performed by CoRAL reconstructs plausible structures based on selecting a minimum number of cycles that are consistent with the constraints provided by long-reads, and together, the cycles explain most of the copy number of the amplicon. On simulated data, most structures were correctly predicted, and even when they were not, they were only slightly rearranged from the true structure. Similarly, in experimental data from cancer cell-lines, the three heaviest reconstructed structures typically explained most of the copy number. In most cases, the reconstruction provides a reasonable template for downstream functional studies, including analysis of regulatory rewiring and chromatin conformation. Of note, CoRAL’s approach can be seamlessly employed for any long-read sequencing technology, such as Oxford Nanopore Technologies or PacBio, where longer reads will always improve breakpoint detection and amplicon reconstruction.

It is important to note, however, that long-reads by themselves are not a panacea, and amplicon reconstruction is different than genome assembly. In diploid genomes, only two haploid structures are possible, and the repeated regions are easily spanned by current long-read technology, except in a few highly repetitive regions. In contrast, larger regions can occur with multiple copies on a single ecDNA, making it hard to resolve into one correct structure. Moreover, heterogeneity of ecDNA may lead to many structures being present. The ecDNA structures resolved by CoRAL may only reflect the most abundant structures. Moreover, due to the minimization of cycle counts, it is possible that the heaviest cycle given by CoRAL glues together smaller ecDNA cycles that share the same segments. To avoid such cases we limit the times that each discordant edge can be traversed in a cycle or walk based on empirical observations. Reconstructions from simulated and real ecDNA amplicons (e.g. COLO320DM) suggested similar cycle sizes to either ground truth or previous characterizations. These considerations will be revised as additional data becomes available.

Thus, we highlight a few avenues for extending and improving CoRAL. First, when a sample has concurrent short-read sequencing data, one may explore if incorporating low-coverage long-reads (< 5X) are sufficient for a hybrid reconstruction. However, due to the rapid evolution of cancer genomes and spatial heterogeneity of tumor samples, the benefit of such an approach may only exist when short and long reads are simultaneously generated from the same biospecimen. Second, CoRAL can be extended to identify the architectures of chromosomal amplicons such as breakage fusion bridge cycles, and ecDNA that have reintegrated into the genome. Because the reconstruction methods use only abstractions relating to path constraints and explained copy number, they can be adapted to other amplifications readily, and this will also be a focus of future studies. Third, as our understanding of amplicon structure grows with experimentally verified structures, that information can be used to improve the constraint space and optimization criteria for CoRAL, and to enhance the simulations of ecDNA or other chromosomal amplifications.

Previous state-of-the-art tools using short-reads like AA (Deshpande et al. 2019) are very accurate in determining if a focal amplification is mediated by ecDNA formation, and in determining the amplified regions. However, they have difficulties in reconstructing the full structure, or determining all regions that participate in one ecDNA molecule. These challenges are partially resolved by targeted deep profiling of a specific subset of amplicons at the expense of not observing the full amplification landscape (Hung et al. 2022). CoRAL not only offers improvements as a standalone tool, but can also be used in conjunction with the targeted approaches - either by refining existing reconstructions or by providing more accurate and unambiguous reconstructions of complex amplicons in targeted enrichment protocols. In summary, CoRAL will be a valuable tool in the arsenal for analyzing complex focal amplifications - such as ecDNA - in tumor genomes, especially as long-read technologies continue to offer cheaper, longer, and more accurate reads.

## Methods

CoRAL takes mapped long-reads (in BAM format) as input, constructs a copy-number-weighted breakpoint graph, decomposes the breakpoint graph into a collection of cycles or paths, and outputs the reconstructed breakpoint graph as well as the resulting cycles/paths from decomposition of the breakpoint graph.

Below, we start with an abstract definition of the breakpoint graph followed by a high-level description of the construction. The copy-number-weighted breakpoint graph (Bafna and Pevzner 1996; Alekseyev and Pevzner 2009; Lin et al. 2014; Deshpande et al. 2019; Hadi et al. 2020), denoted by 𝒢 = (*V, E* = *E*_*s*_ ∪ *E*_*c*_ ∪ *E*_*d*_, CN), encodes a collection of non-overlapping intervals on a given reference genome, which are amplified, reordered or reoriented. A brief description is provided here with details in Appendix A1:

- Each *v ∈ V* represents the start or end coordinate of an interval, or the special *source* nodes *s, t* (defined below). Let *l*_*v*_ denote the location of node *v*.
- *E*_*s*_ represents *sequence edges*, that join the start and end coordinates of an interval.
- *E*_*c*_ represents *concordant edges* so that (*u, v*) *∈ E*_*c*_ if *l*_*v*_ − *l*_*u*_ = 1 where *v* is the start coordinate of the canonically larger interval on the reference genome represented by a sequence edge.
- *E*_*d*_ represents *discordant edges*, generated when (sufficient) reads map to discordant intervals. Thus, (*u, v*) *∈ E*_*d*_ if |*l*_*v*_ − *l*_*u*_| ≠ 1, if the read connecting *u* to *v* changes orientation, or if the nodes are on different chromosomes. A discordant edge could connect the start (or end) coordinate of an interval to itself (an inverted duplication or *foldback* ).
- All edges are weighted using the real-valued function CN : *E* → ℚ+ denoting the *copy number*. The CN is computed based on an assumption of diploidy for the majority of basepairs on the genome. We require that the CN assignment be “balanced”, for each (*u, v*) *∈ E*_*s*_, as follows:

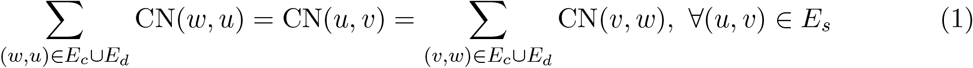

By definition, each node in a breakpoint graph is connected to a single sequence edge, and a single concordant edge as well; but it may connect to multiple discordant edges. The source nodes *s* and *t* connect to the canonically smallest and canonically largest coordinate on the reference genome from a collection of consecutive intervals connected by concordant edges; or a sequence edge which is only connected to another sequence edge with smaller CN by concordant edges, and therefore is deemed to violate the balanced CN constraint without the source connections. Edges connected to source nodes are treated as discordant edges. See 1b for an example of a breakpoint graph constructed by CoRAL from the ecDNA in 1a. We denote a maximal collection of genomic intervals connected by concordant edges as an *amplified interval*, and the union of all amplified intervals and their (discordant) connections as an *amplicon*. Note that a tumor sample could contain multiple amplicons whose intervals are non-intersecting. CoRAL constructs a distinct breakpoint graph for each amplicon.

### Breakpoint graph construction with CoRAL

To build the breakpoint graph for an amplicon, CoRAL first determines all amplified intervals included in the amplicon. CoRAL requires *seed amplified intervals* (Supplementary Figure 11a) as a starting point to search for all connected amplified intervals contained in an amplicon. The seed amplified intervals can be derived from whole genome CNV calls (e.g., with third party tools like CNVkit (Talevich et al. 2016)) of mapped long reads. From the CNV calls, we select the genomic segments adjacent to each other with a minimum threshold of copy number as well as the aggregated size as seed intervals (Appendix A2).

With the seed amplified intervals, CoRAL searches for amplified intervals connected to seed intervals (by discordant edges) using a breadth first search (BFS). For BFS CoRAL maintains a list ℐ of amplified intervals in all amplicons it explored or discovered so far, initialized as the list of seed intervals; and a set ℰ representing the connections between amplified intervals through discordant edges. Each pair of intervals (*a*_*i*_, *a*_*j*_) *∈ ℰ* (*i* ≤ *j*) is labeled by the breakpoints connecting two loci within the intervals *a*_*i*_ and *a*_*j*_ respectively. The main iteration explores the next unvisited interval *a*_*i*_ in ℐ, indicating a new amplicon (connected component), until all amplified intervals in ℐ are visited.

Let *L* be a priority queue used in the interval search starting from *a*_*i*_, which is initialized with a single element *a*_*i*_. Each step of the interval search pops the first interval *a*_0_ in *L*, and extracts all breakpoints supported by chimreic alignments connecting a locus within *a*_0_ to another locus on the reference genome. These breakpoints are greedily clustered (with the procedure described below) and the new locus *l* determined by a cluster 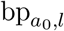 of breakpoints of size at least haploid coverage is chosen to be further explored (Supplementary Figure 11b). If the new locus falls into an existing interval *a*_*e*_ *∈* ℐ, then mark interval *a*_*e*_ as visited, augment the label set of (*a*_0_, *a*_*e*_) with 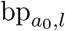, and only append *a*_*e*_ to *L* if it was not previously visited. Otherwise CoRAL will extend *l* to a new amplified interval including *l*, depending on whether *l* is amplified from the CNV calls. If *l* is amplified, CoRAL will append the new interval *a*_*n*_ = [chr_*l*_, max(*s*_*l*_ − *δ, l* − Δ), min(*e*_*l*_ + *δ, l* + Δ)] to both *L* and *I*, where *s*_*l*_ and *e*_*l*_ are the start and end coordinate of the amplified CN segments including *l* in CNV calls. If *l* is not amplified, CoRAL will append the new interval *a*_*n*_ = [chr_*l*_, *l* − *δ, l* + *δ*] to *L* and *I*. In either case, CoRAL also labels the connection (*a*_0_, *a*_*n*_) with 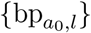 and add it to ℰ. The amplified interval search starting from *a*_*i*_ is repeated until *L* becomes empty. A pseudocode of the above procedure as well as the selection of Δ and *δ* is discussed in detail in Appendix A2.

At the end of interval search, all intervals *I* are visited, and each connected component of amplified intervals by breakpoint edges with sufficient support of long reads forms an amplicon (Supplementary Figure 11c). After BFS, CoRAL postprocesses the amplified intervals discovered in *I* by merging (i) adjacent (in CNV calls) or overlapping intervals, or (ii) intervals on the same chromosome which are not adjacent but have close (i.e., within ≤ 2*δ*-bp vicinity) breakpoint connections. Two intervals belonging to different amplicons are brought into the same amplicon after merging. CoRAL will then search for breakpoints within a single (merged) amplified interval (Supplementary Figure 11d). Finally, CoRAL builds the actual breakpoint graph for each amplicon. It will split each amplification interval into sequence edges if there are breakpoint edges connecting to the middle of that interval, and add concordant edges connecting two adjacent sequence edges on the reference genome (Supplementary Figure 11e).

### CN assignment

Once the graph structure 𝒢 is fixed, CoRAL recomputes the CN for each edge in 𝒢 (Supplementary Figure 11e), as the initial CNV calls used for amplified interval search may not follow the balance requirement (Eqn. 1), and they do not account for concordant and discordant edges. Let the diploid long read coverage be *θ*. CoRAL assumes that the majority of the donor genome is not amplified and estimates *θ* as the coverage on the 40-th percentile of CN segments sorted by their initial CNV calls. Given *θ*, CoRAL models the total number of nucleotides on each sequence edge (*u, v*) *∈ E*_*s*_ as a normal distribution with mean and variance *θ* · CN(*u, v*) · *l*(*u, v*), where *l*(*u, v*) denotes the length (in bp) of the sequence edge; and the number of reads supporting each concordant and discordant edge (*u, v*) *∈ E*_*c*_ ∪ *E*_*d*_ as a Poisson with mean *θ* · CN(*u, v*). To estimate CN, CoRAL computes the maximum likelihood of CN using the joint distribution of observed number of nucleotides on each sequence edge and the observed read counts on each concordant/discordant edge–with the constraint that CN is balanced (Appendix A3). The (convex) optimization problem was solved using CVXOPT package.

### Cycle extraction

We are interested in paths and cycles that alternate between sequence and breakpoint (i.e., concordant or discordant) edges, thus by definition, if the path contains node *s* (respectively, *t*), it must be the first (respectively, last) node in the path. Define an *alternating sequence* of nodes as a sequence *v*_1_, *v*_2_, …, *v*_*w*_, where for all 1 ≤ *i* < *w*, (*v*_*i*_, *v*_*i*+1_) *∈ E*, and the edges alternate between sequence and breakpoint edges. Define a *walk* in *G* as an alternating sequence *v*_1_, *v*_2_, …, *v*_*w*_, where *v*_1_ = *s, v*_*w*_ = *t*. A *path* is a walk with no node repeated (*v*_*i*_ = *v*_*j*_ ⇔ *i* = *j*). A *cyclic walk* or *cycle* is an alternating sequence *v*_1_, *v*_2_, …, *v*_*w*_ of nodes where *v*_1_ = *v*_*w*_ *≠ s, t*. The cycle is simple if no node except the first/last one is repeated.

The amplicon encoded by 𝒢 is composed of a superposition of cycles and walks with high copy numbers. For all sequence edges (*u, v*) *∈ E*_*s*_, define the *length-weighted copy number* using C_*l*_(*u, v*) = CN(*u, v*) · *l*(*u, v*). Similarly, for graph 𝒢.

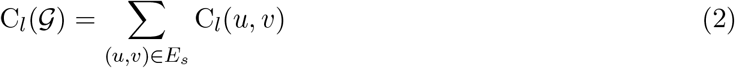

Our goal is to identify a minimum number of cycles and walks (denoted as *W*_*i*_), each associated with a positive weight – corresponding to the copy number (based on the assumption of uniform coverage, see Supplementary Figure 12) – so that the sum of weights on all edges in all walks composes a large fraction of C_*l*_(𝒢). Furthermore, the long-reads that span multiple (at least 2) breakpoints in 𝒢, also provide us with a collection of subwalks 𝒫 = *{p*_1_, …, *p*_*m*_*}*, and the reconstructed walks must simultaneously be consistent with a large fraction of these subwalks (Appendix A4).

The complexity of cycle extraction and rationale for CoRAL’s optimization procedure can be motiviated through an example illustration (Supplementary Fig. 13). The breakpoint graph in Supplementary Fig. 13a consists of segments A, B, and C assumed for simplicity to be of equal length. The optimization in CoRAL will decompose it into a single cycle of copy number 50, with a duplication of segment B (right panel). The decomposition is also supported by the subwalk constraint given by the long read that connects segment A, B and C. Even if the long-read were not present, this cycle is still a parsimonious solution compared to an alternative decomposition with two cycles (one containing A and B, and the other containing B and C).

Supplementary Figure 13b has a similar graph structure but with different copy numbers on segments. The best decomposition is given by one cycle containing A and B with copy number 80, and a second cycle containing A, B(2 copies), C with copy number 10. The total length weighted copy number of the graph is 200 (assuming segments of length 1). Cycle 1 explains 80% and Cycle 2 explains 20% of the copy number. Other decompositions of cycles are indeed possible. For example, if the subwalk constraint given by the long read were not present, an alternative decomposition with 90 copies of Cycle 1 and 10 copies of a different cycle 2 containing only segments B and C would also be explain all length-weighted copy numbers in the graph. On the other hand, a more parsimonious decomposition into one single cycle with copy number 10, where segment A repeating 9 times and segment B repeating 10 times is not allowed because it violates the upper bound on the multiplicity of segments in the cycle (see auxiliary constraint 3 in the MIQCP formulation below). For the same reason, the decomposition into one single cycle is not allowed in Supplementary Figure 13c.

We resolve the multi-objective challenge using mixed integer quadratically constrained programming (MIQCP). The MIQCP works with 2 parameters: *α* as the minimum fraction of length-weighted copy number explained, and *β* as the minimum fraction of path constraints satisfied. Additionally, parameter *k* denoting the maximum number of cycles/walks allowed is learned starting with *k* = 10, according to two modes. In the **full** mode (MIQCP-full) desscribed below, the MIQCP attempts a solution with at most *k* walks that satisfy other constraints, or returns ‘infeasible’. The value of *k* is doubled until feasibility is reached or *k* > |*E*|. The **greedy mode** is described later. We implement both quadratic programs with through the python3 interface of Gurobi 10.0.1.

MIQCP-full utilizes the following **key** variables.

- *w*_*i*_ *∈* ℚ ≥ 0: denotes the copy number for walk *W*_*i*_ (1 ≤ *i* ≤ *k*); and *z*_*i*_ *∈ {*0, 1*}* indicates if *w*_*i*_ > 0;
- *x*_*uvi*_ *∈* ℤ ≥ 0 represents the number of times walk *W*_*i*_ traverses (*u, v*) for each edge (*u, v*) *∈ E* and 1 ≤ *i* ≤ *k*;
- *P*_*j*_ *∈ {*0, 1*}* indicates if subwalk constraint *p*_*j*_ is satisfied for 1 ≤ *j* ≤ *m* ;

The MIQCP(*k, α, β*) objective is given by:

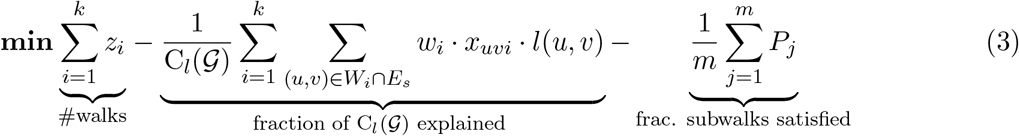

subject to the constraints:

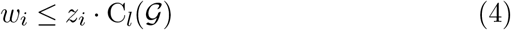

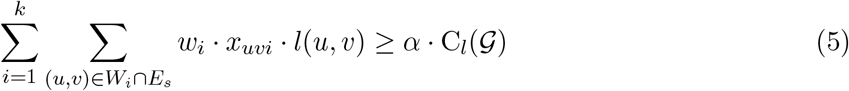

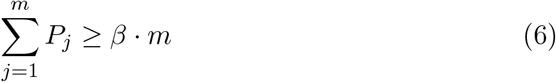

Constraint 4 ensures that *w*_*i*_ = 0 if *z*_*i*_ = 0 ∀*i* = 1, …, *k*, and constraints 5,6 ensure that minimum fractions of the length weighted copy number and subwalk constraints are satisfied. The unsatisfied fractions also contribute a small amount to the MIQCP objective. To ensure that cycles and walks have their nodes connected, alternating-edge structure, we must satisfy several auxiliary constraints, enumerated below, with details in the Appendix (Equations A4.6-A4.23).

1. Each *W*_*i*_ should form a valid walk of alternating sequence and breakpoint (i.e., concordant or discordant) edges.
2. The total CN of all cycles/walks passing through an edge (*u, v*) *∈ E* is at most C_*l*_(*u, v*).
3. We require that each cycle/walk traverses through a discordant edge (*u, v*) at most *R*(*u, v*) times. By default the value of *R*(*u, v*) is estimated for each discordant edge (*u, v*) *∈ E*_*d*_ based on the number of (long) reads supporting that edge. See Appendix A4 for details.
4. Each walk *W*_*i*_ (if *z*_*i*_ > 0), either forms a cycle starting at node *v*_1_*≠ s*, or starts at *s* and ends at *t*. If *W*_*i*_ forms a cycle we require that the concordant or discordant edge connected to *v*_1_ occurs only once in the cycle.
5. *x*_*uvi*_ and *z*_*i*_ are consistent. *z*_*i*_ = 1 ⇔ *x*_*uvi*_ > 0 for some (*u, v*) *∈ E*.
6. **Connectivity**. We use auxiliary variables to encode the “discovery order” of the nodes in walk *W*_*i*_. These variables number the nodes from ‘1’ for the start node, and incrementing by one for each susbequent node in the cycle/walk.
7. **Subwalk constraints**. We enforce a weak constraint by requiring each walk *p*_*j*_ *∈ 𝒫* to be present as a subgraph of the graph induced by some walk *W*_*i*_.

### MIQCP-greedy(*α, β, γ, ϵ*)

For a large graph (e.g. |*E*| > 100), MIQCP-full could be resource intensive. Therefore, we also implemented an MIQCP with additional parameters *γ, ϵ*, but not *k*, that identifies only a single walk maximizing the copy number, and additional subwalk constraints satisfied, with parameter *γ* controlling the weight of the two objectives. Let

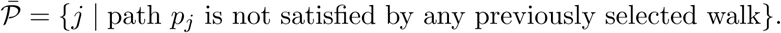

Then the greedy MIQCP objective to identify the next walk *W*_*i*_ is given by:

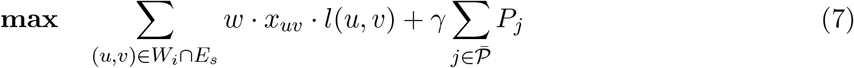

Each time a new walk is computed, its copy number is removed for all edges it passed through, and 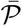 is updated. The procedure is repeated until either *α* · C_*l*_(𝒢) copy numbers and *β* · *m* subwalk constraints are explained by the currently selected walks, or the copy number of next walk is less than *ϵ* · C_*l*_(𝒢), for parameter *ϵ*. We empirically set 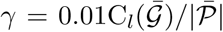, where 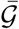 denote the remaining length-weighted copy number of 𝒢 after removing the copy numbers from the last walk, and *ϵ* = 0.005. The greedy MIQCP is solved using the same set of auxiliary constraints as before.

### Implementation details

In practice, if *G* has |*E*| > 100 edges, we use the iterative greedy MIQCP, until either 90% of CN weight is removed from the graph, or the CN weight of the next cycle is less than 1% of the total CN weight. Otherwise, we run full-MIQCP with *α* = 0.9, *β* = 0.9. Initially, *k* = 10, and it is doubled until a feasible solution is reached. If doubling the number of cycles/paths leads to more than 10000 variables in the integer program, we switch to greedy-MIQCP. CoRAL provides users an option to postprocess the greedy-MIQCP solutions with full MIQCP with 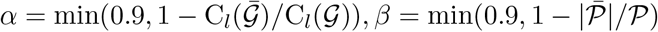.

## Data Access

All raw and processed sequencing data generated in this study have been deposited to the NCBI BioProject database under accession number PRJNA1110283 (https://www.ncbi.nlm.nih.gov/bioproject/PRJNA1110283). We also obtained short read and long read sequencing data for COLO320-DM, COLO320-HSR, GBM39, GBM39-HSR and CHP-212 cells from the NCBI Sequence Read Archive under BioProject accessions PRJNA506071, PRJNA670737 and PRJNA622577.

### Code availability

The version of CoRAL used in the presented analysis is included as Supplemental Code. An up-to-date version of CoRAL is available on GitHub https://github.com/AmpliconSuite/CoRAL.

## Acknowledgements

We thank members of the Bafna, Chang, and Mischel labs as well as members of eDyNAmiC for helpful discussion and feedback. We thank Mădălina Giurgiu and the Henssen lab for helpful discussion around running and interpreting results from Decoil. We thank Suhas Srinivasan for technical assistance in cloud computing.

This work was delivered as part of the eDyNAmiC team supported by the Cancer Grand Challenges partnership funded by Cancer Research UK (CGCATF-2021/100012 [P.S.M., H.Y.C.]; CGCATF-2021/100025 [V.B.]) and the National Cancer Institute (OT2CA278688 [P.S.M., H.Y.C.]; OT2CA278635 [V.B.]); NIH R35-CA209919 (H.Y.C.); U24CA264379, R01GM114362 (V.B.); and National Cancer Institute of the National Institutes of Health under Award Number K99CA286968 (M.G.J.). K.L.H. was supported by a Stanford Graduate Fellowship and an NCI Predoctoral to Postdoctoral Fellow Transition Award (NIH F99CA274692). H.Y. is a Howard Hughes Medical Institute Awardee of the Life Sciences Research Foundation. H.Y.C. is an Investigator of the Howard Hughes Medical Institute. The content is solely the responsibility of the authors and does not necessarily represent the official views of the National Institutes of Health.

## Author Contributions

K.Z., M.G.J., J.L., and V.B. conceived the project. K.Z. and V.B. conceived of the CoRAL algorithm with input from M.G.J. and J.L. and assumptions and optimizations. K.Z. developed the CoRAL software with assistance from M.G.J. and J.L. H.Y., I.T.L.W., and S.Z. provided monoclonalized cell lines. K.L.H. performed Nanopore sequencing data for GBM39 and GBM39-HSR cell lines. M.G.J. performed Nanopore and Illumina sequencing GBM39 (mono), COLO320-DM (mono), PC3-DM (mono), and PC3-HSR (mono) cell lines. M.G.J. performed all simulations using ecSimulator and implemented benchmarking pipelines for AmpliconArchitect, CoRAL, and Decoil, with input from J.L. and K.Z. K.Z. performed reconstruction of amplicons using CoRAL and AmpliconArchitect on all cell line data and analyzed results. X.B., K.Z., and J.L. developed software for visualizing reconstructed amplicons. V.B., P.S.M., and H.Y.C. guided data analysis, provided feedback on experimental design, and supervised this work. K.Z., M.G.J., J.L., and V.B. wrote the manuscript with input from all authors.

## Supplementary Figures

**Supplementary Figure 1:**
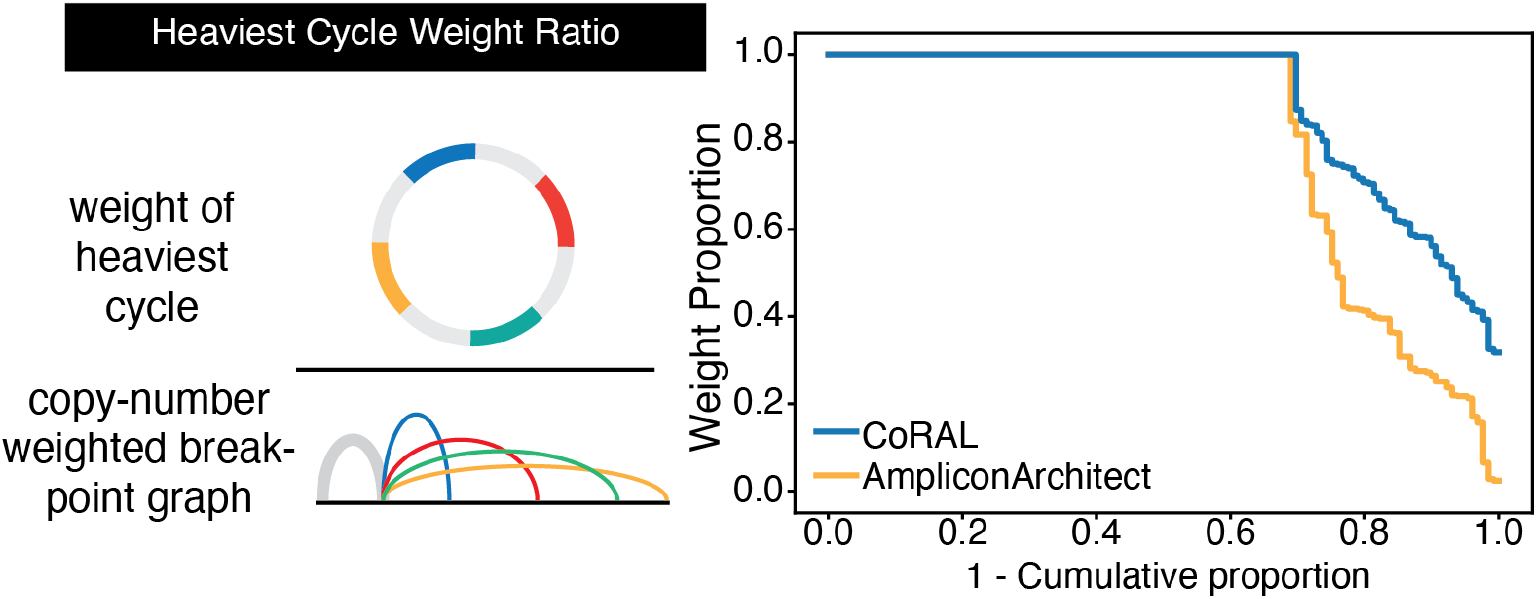
Fraction of length-weighted copy numbers given by the heaviest cycle from AA and CoRAL reconstructions. Fraction of length-weighted copy numbers given by the heaviest cycle over the total length-weighted copy numbers in the inferred breakpoint graph (i.e., “Heaviest cycle weight ratio” below) is reported across all simulated amplicons.

**Supplementary Figure 2:**
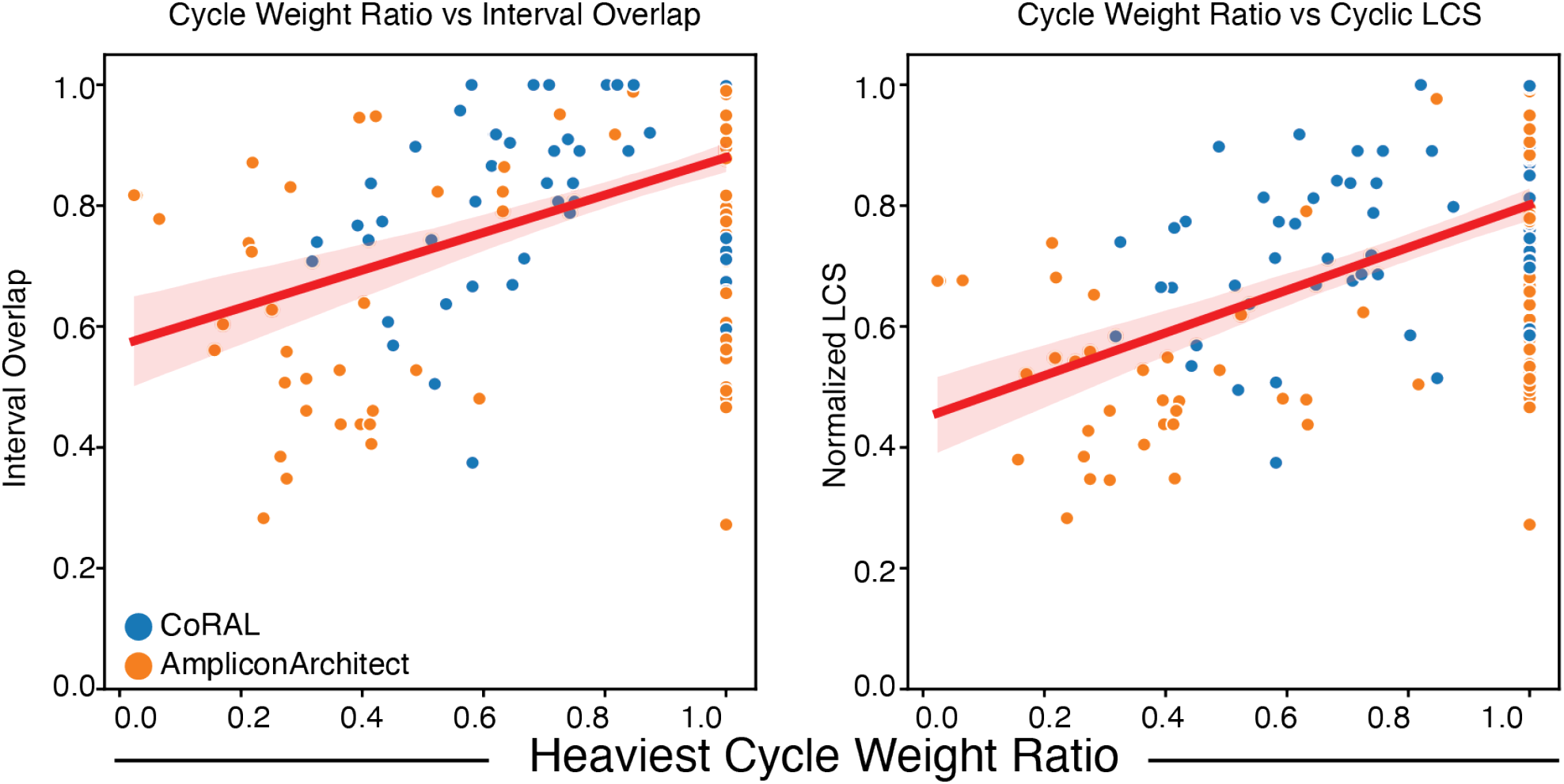
Comparison of the cycle weight ratio vs accuracy measures of simulated benchmarks.

**Supplementary Figure 3:**
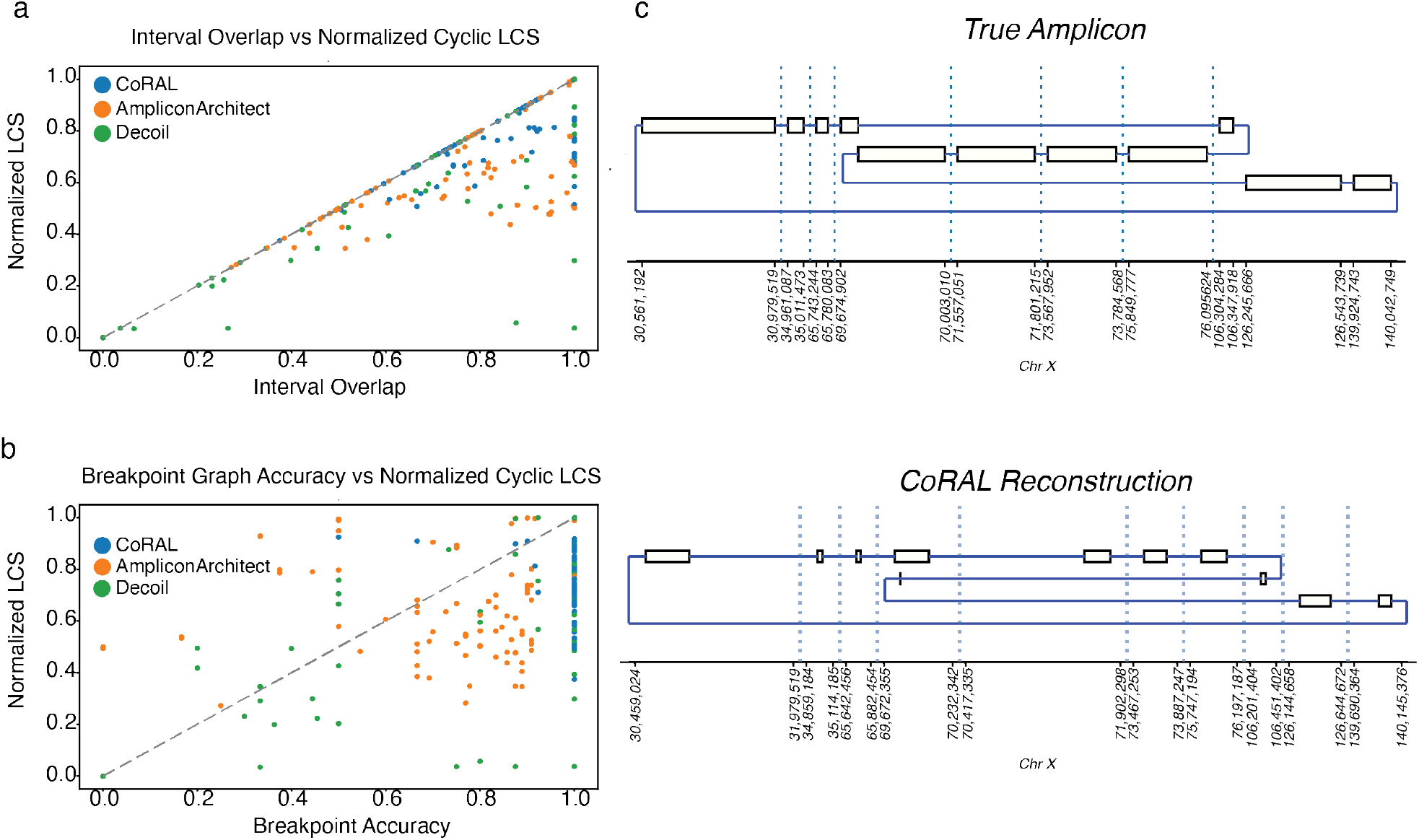
Reconstructions with perfect overlap but imperfect ordering. Evaluation of the Normalized Cyclic LCS (which captures ordering of intervals) vs the interval overlap (a) and breakpoint accuracy (b). An example CoRAL reconstruction with an interval overlap of 0.99 but a normalized cyclic LCS of 0.51 due to an incorrectly inverted set of intervals.

**Supplementary Figure 4:**
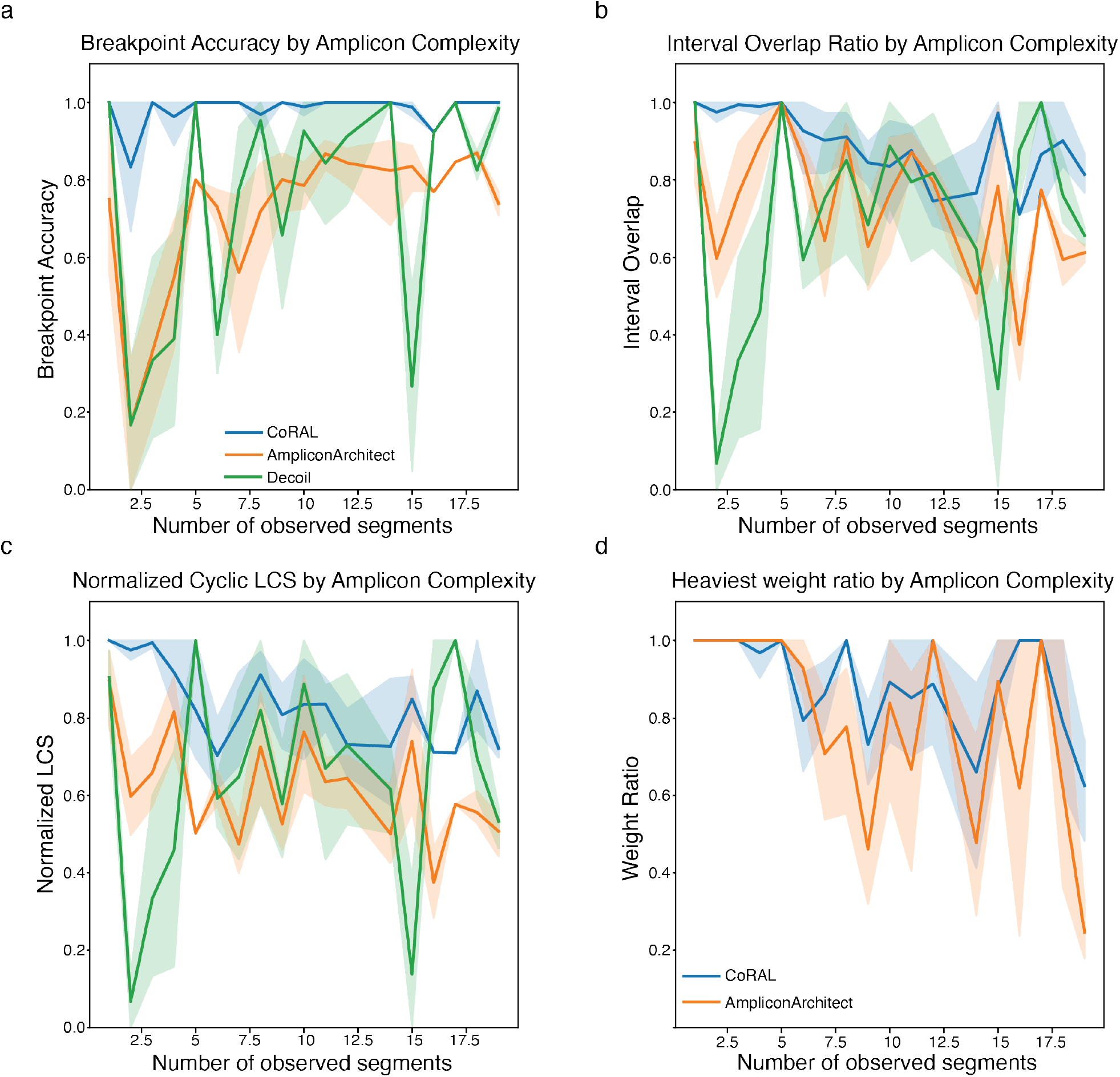
Performance on simulated data separated by number of genome segments in the true amplicon. Performance of CoRAL, AmpliconArchitect, and Decoil (a-c only) on simulated amplicons as a function of the number of simulated genome segments, capturing breakpoint accuracy (a), interval overlap ratio (b), normalized cyclic longest-common substring (LCS; c), and heaviest cycle weight ratio (d).

**Supplementary Figure 5:**
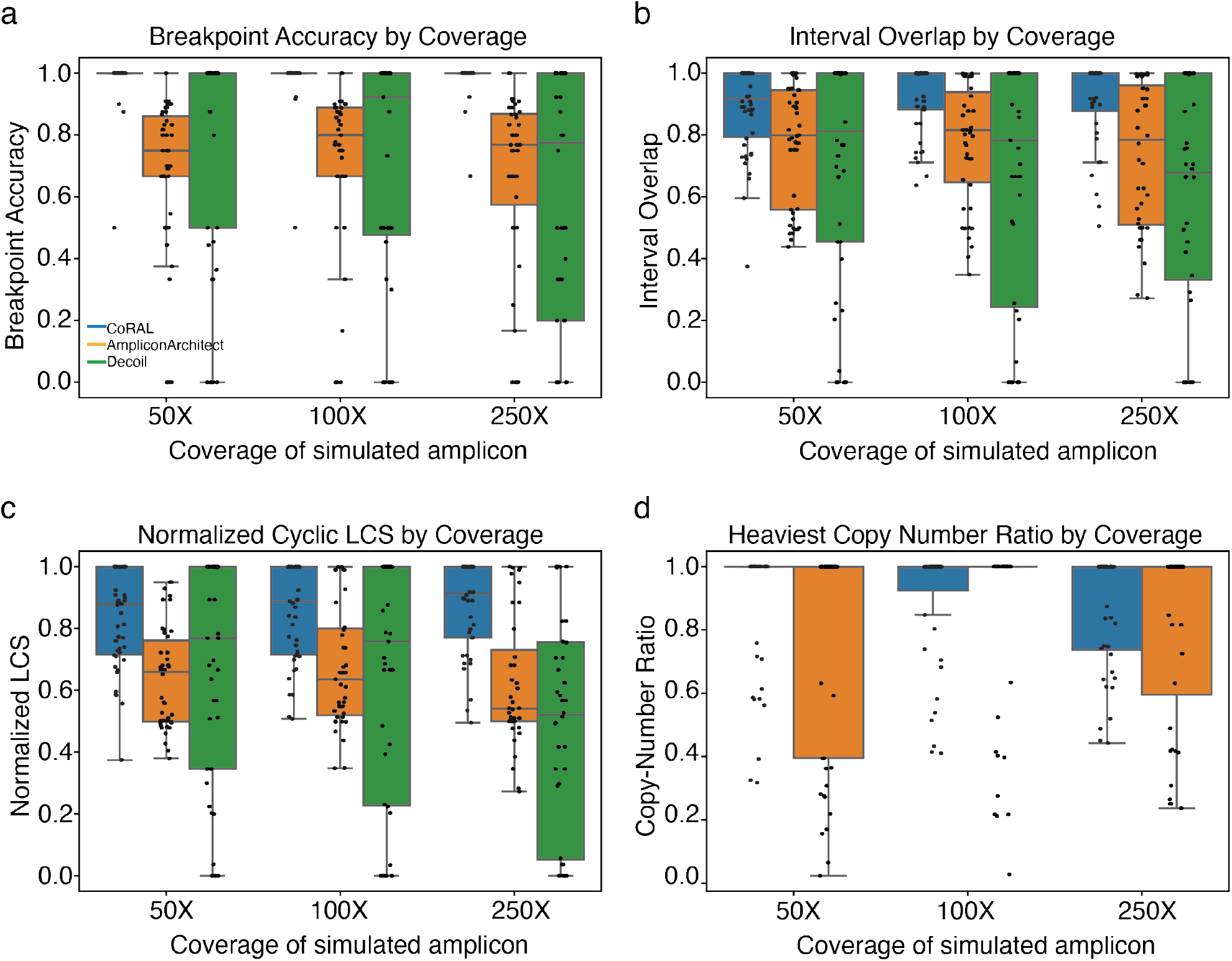
Accuracy of simulations by simulated amplicon coverage. Break-point accuracy (a), interval overlap ratio (b), normalized cyclic longest-common substring (LCS; c), and heaviest cycle weight ratio (d) for simulations broken down by simulated amplicon coverage.

**Supplementary Figure 6:**
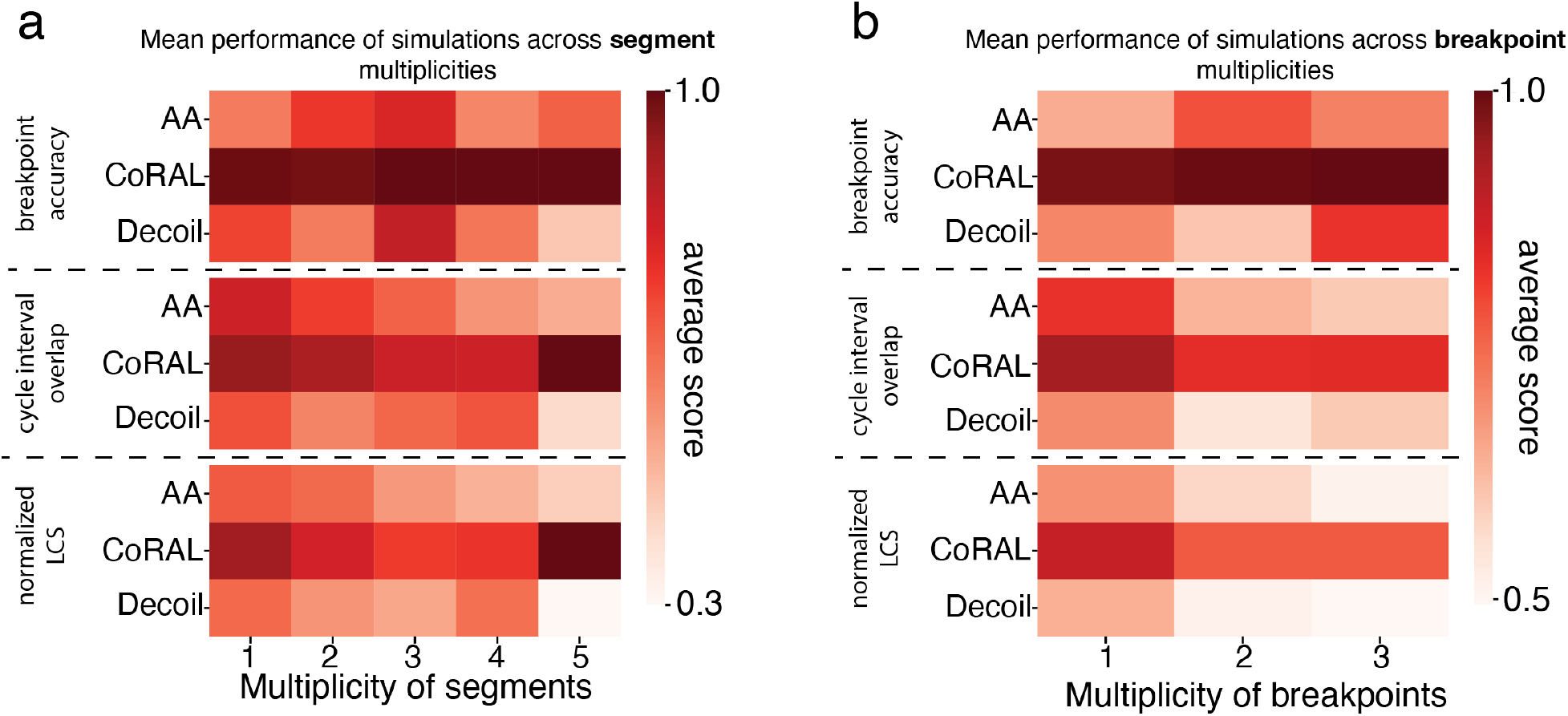
Accuracy of simulations by multiplicity of segments and break-points in amplicon. Breakpoint accuracy, interval overlap ratio, and normalized cyclic longest-common substring (LCS) for simulations reported as a function of (a) largest multiplicity of segments, and (b) largest multiplicity of breakpoints. Values in heatmaps correspond to the mean performance across replicates.

**Supplementary Figure 7:**
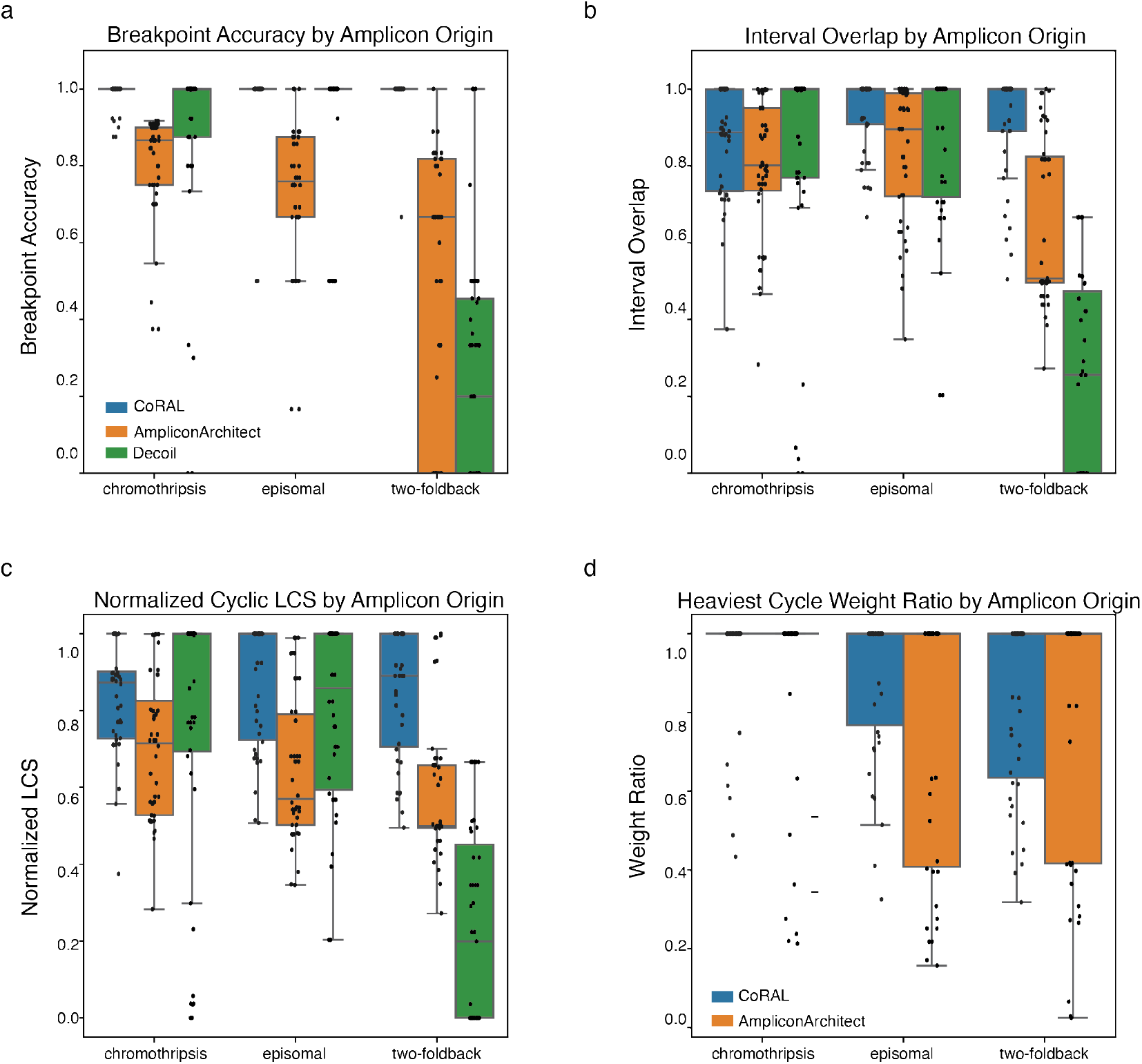
Accuracy of simulations by simulated amplicon origin. Breakpoint accuracy (a), interval overlap ratio (b), normalized cyclic longest-common substring (LCS; c), and heaviest cycle weight ratio (d) for simulations broken down by the origin mechanism of simulated amplicon.

**Supplementary Figure 8:**
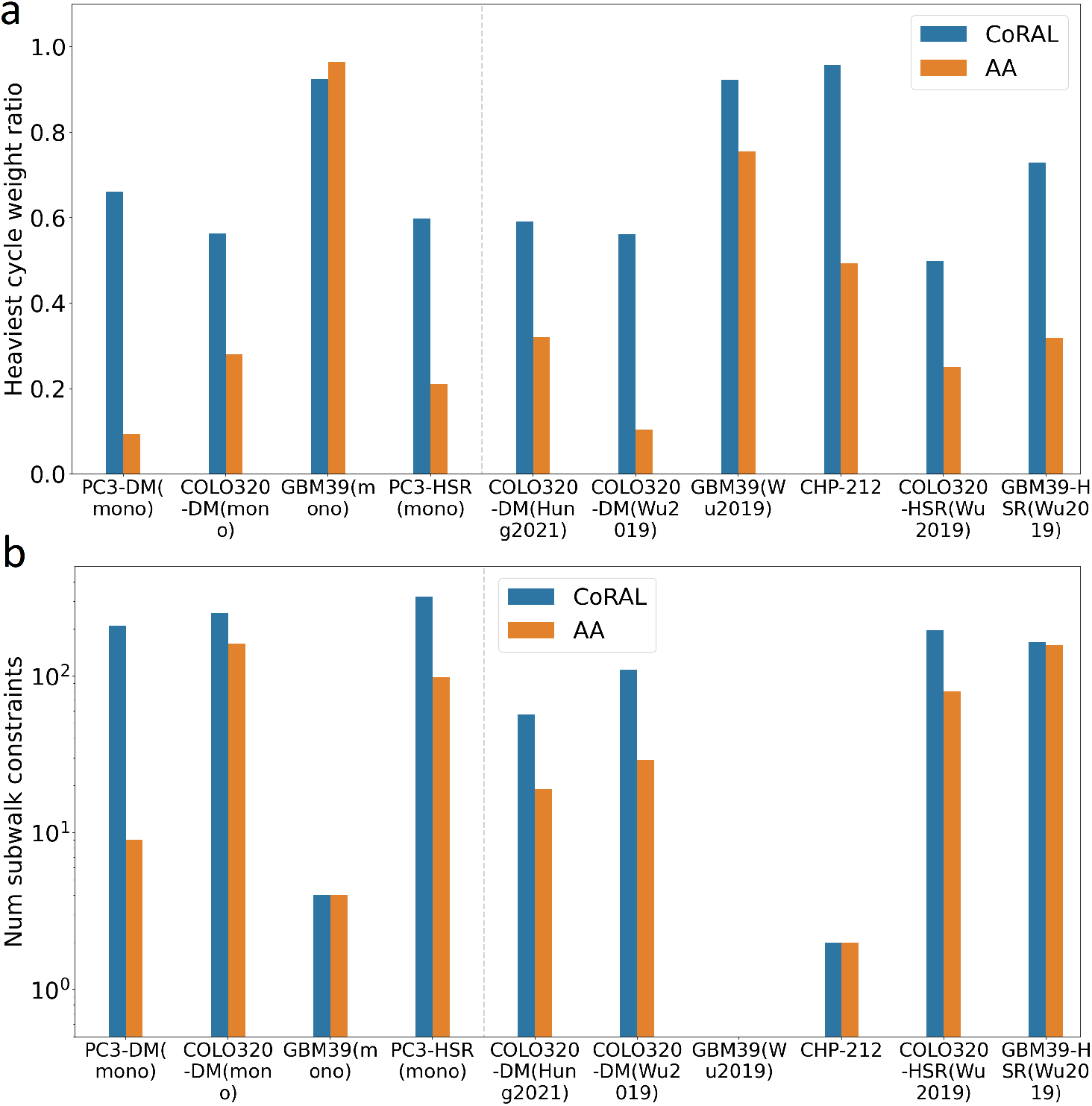
Amplicon reconstruction performance in cell lines, with a single heaviest cycle. (a) Heaviest cycle weight ratio from a single heaviest cycle (i.e., *k* = 1) reported by CoRAL and AA; (b) number of subwalk constraints satisfied by the heaviest cycle reported by CoRAL and AA, in cell lines.

**Supplementary Figure 9:**
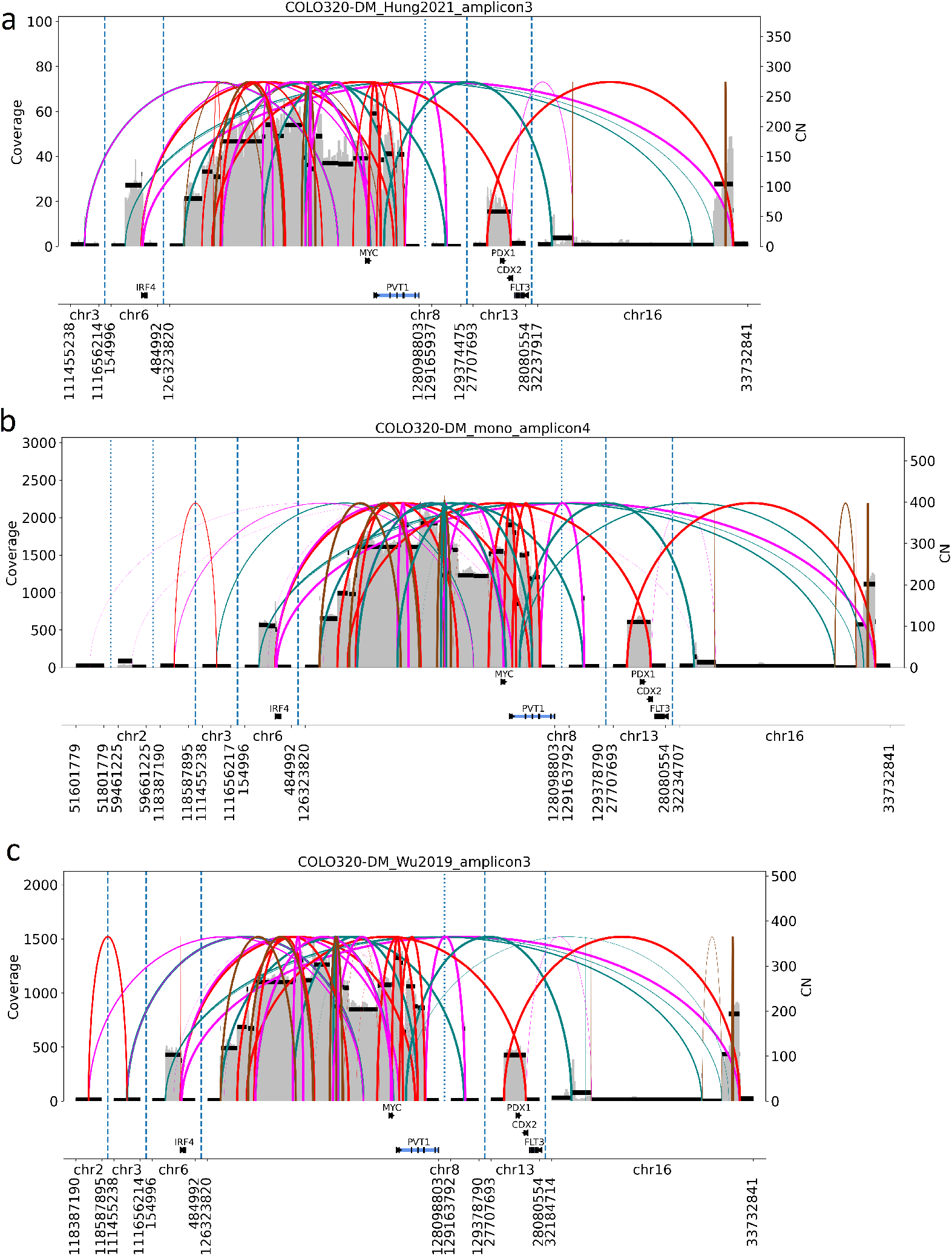
Breakpoint graph reconstructed by CoRAL from the three COLO320-DM samples. (a) COLO320-DM (Hung 2021) breakpoint graph reconstructed by CoRAL. (b) COLO320-DM (mono) breakpoint graph reconstructed by CoRAL. (c) COLO320-DM (Wu 2019) breakpoint graph reconstructed by CoRAL.

**Supplementary Figure 10:**
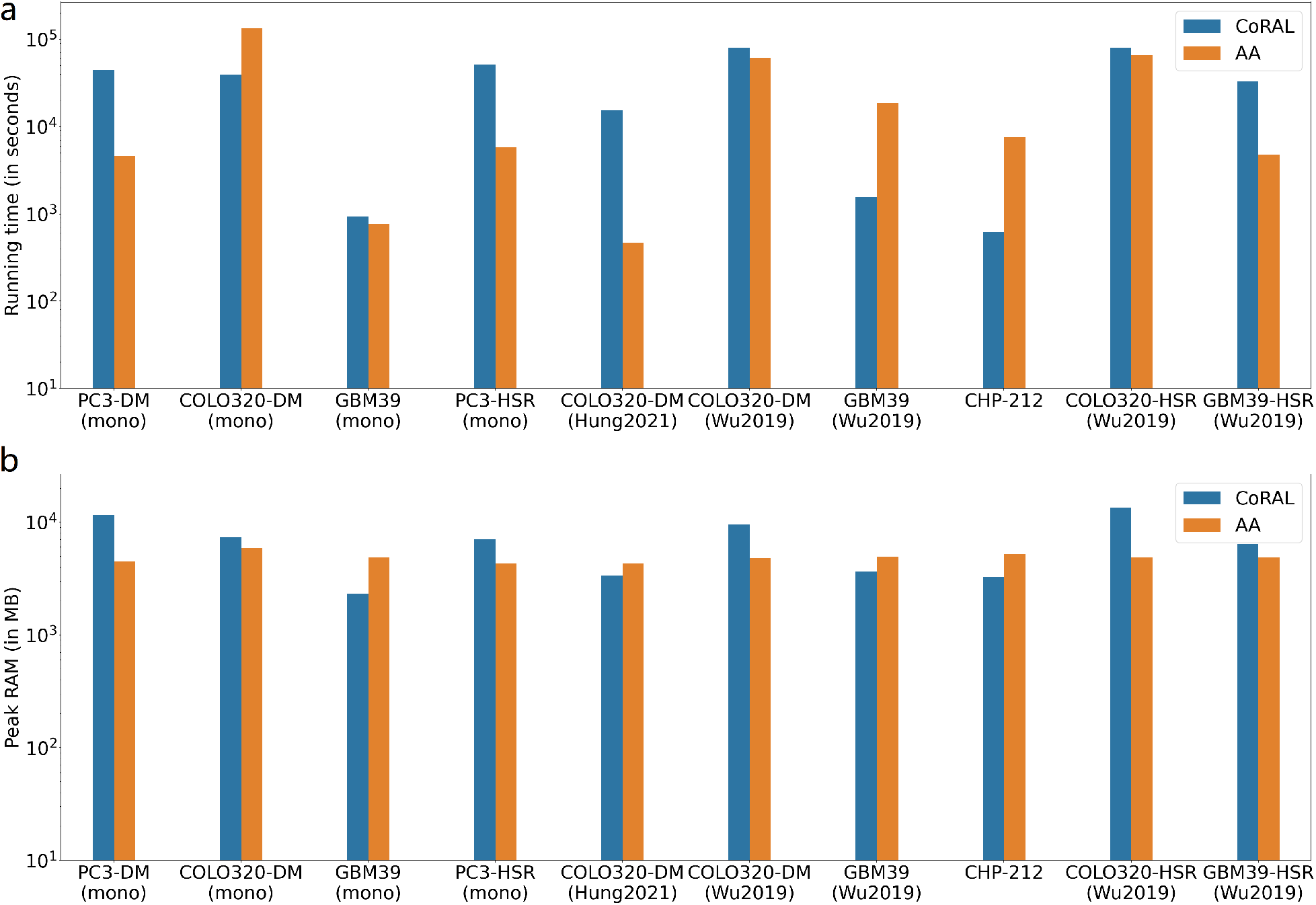
Running time and memory comparison between CoRAL and AA. (a) Running time, in seconds; (b) Peak RAM usage, in Megabytes, to reconstruct all amplicons (and cycles) in each sample.

**Supplementary Figure 11:**
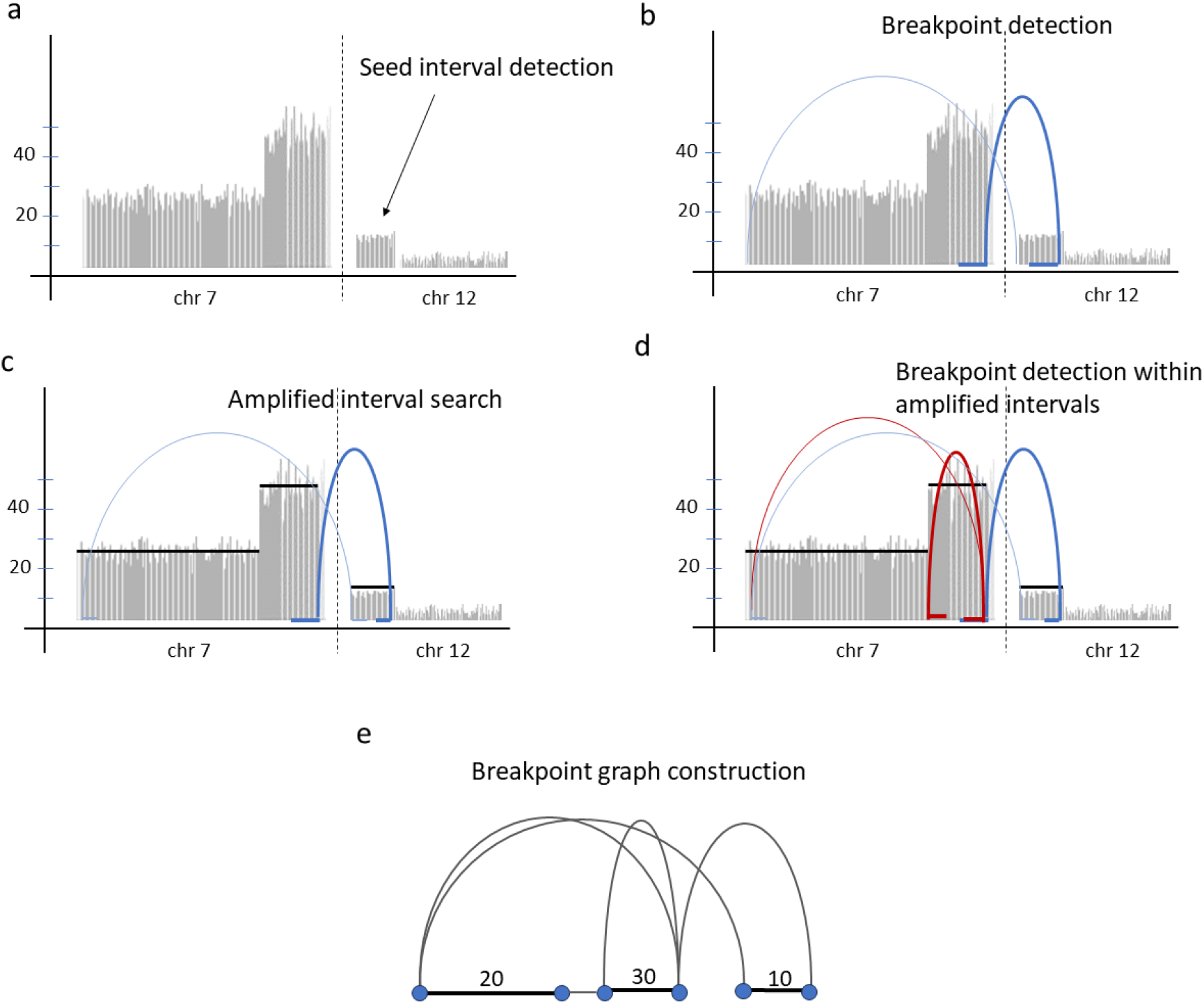
Illustration of amplified interval search and breakpoint graph construction. (a) CoRAL starts with seed amplified intervals to search for all connected amplified intervals contained in an amplicon. (b) CoRAL explores new amplified intervals connected to seed intervals through breakpoints. (c, d) CoRAL first searches for all amplified intervals constituting an amplicon and then detects additional breakpoints within each amplified interval. (e) Finally, CoRAL builds a breakpoint graph with the list of amplified intervals, splitting the intervals at breakpoint junctions.

**Supplementary Figure 12:**
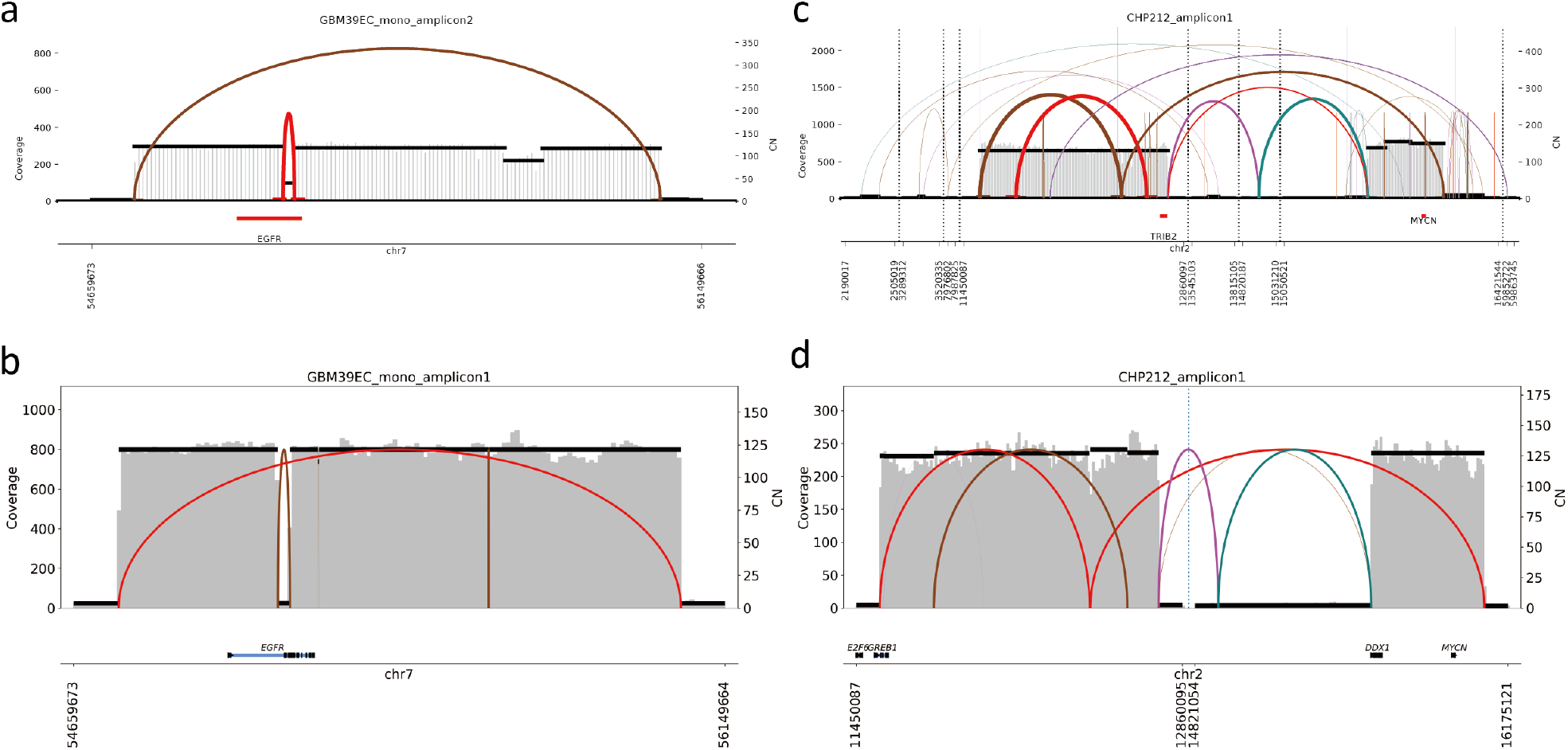
Breakpoint graph reconstructed from cell lines indicating uniform coverage on ecDNA. Breakpoint graph reconstructed from (a) GBM39 (mono) short reads; (b) GBM39 (mono) long reads; (c) CHP-212 short reads; (d) CHP-212 long reads. Gray vertical bars indicate (short/long reads) coverage and black horizontal lines indicate predicted CN for each sequence edge in the graph.

**Supplementary Figure 13:**
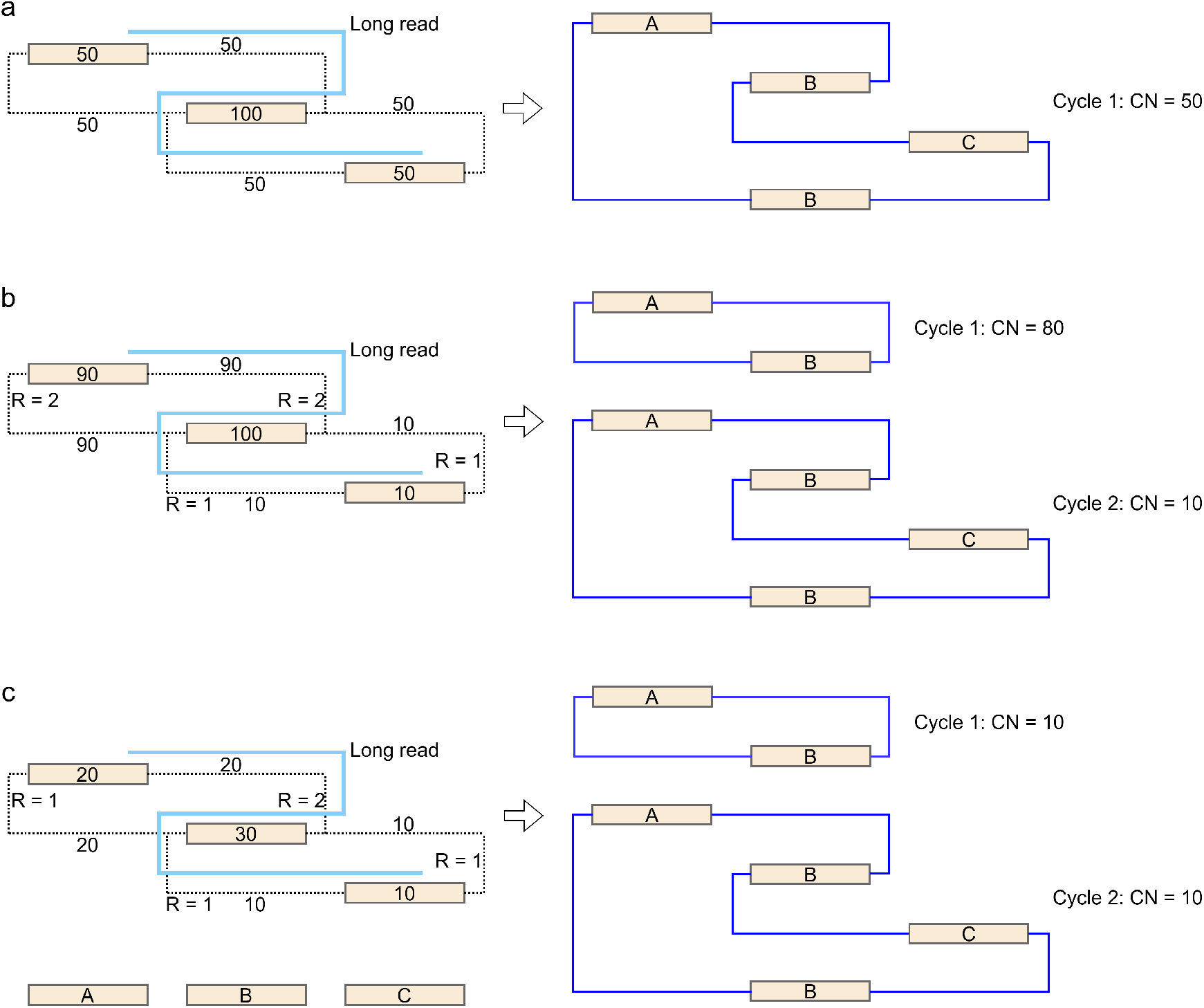
Cycle decomposition with subwalk constraints. (a) A single cycle with copy number 50 containing 2 copies of segment B and 1 each of segments A and C accounts for 100% of the copy number in the breakpoint graph. It also satisfies the subwalk constraint given by the long read. (b) The copy number of the breakpoint graph is explained by a minimum of 2 cycles, with copy numbers 80 and 10 respectively, due to the number of times each discordant edge can be traversed in a cycle (marked on the discordant edge as *R* = 1 or *R* = 2). (c) The copy number of the breakpoint graph is explained by a minimum of 2 cycles, with copy numbers 10 and 10 respectively, again due to a limit on the number of times each discordant edge can be traversed in a cycle.

## Appendix

### A1. Preliminaries

We begin by offering a set of definitions that are used throughout this paper.

#### Reference Genome

A **reference genome** is a set of *R* strings ℛ𝒢 = *{s*_1_, …, *s*_*R*_*}* representing the chromosomes in a genome. For example, the human genome ℛ𝒢 is comprised of *R* = 24 strings typically labeled as *chr*1, …, *chr*22, *chrX*, and *chrY* . We denote the length of string *s*_*i*_ with |*s*_*i*_|. At times we call strings in ℛ 𝒢 **chromosomes** and enforce a total order ≺ on the chromosomes (e.g., the human genome string *chr*2 is smaller than *chr*3 and thus appears before it). A **genomic interval** *s*_*i*_[*l, r*] is a sequence of nucleotides in string *s*_*i*_ starting at position *l* and ending at position *r* (both inclusive) where 1 ≤ *l* ≤ *r* ≤ |*s*_*i*_|. At times, we use *s*_*i*_[*l*, …] to denote an interval starting at position *l* and *s*_*i*_[…, *r*] as a interval ending at position *r*.

#### Breakpoint

A **breakpoint** describes a junction between two genomic intervals in the reference genome set. We describe a breakpoint bp by a pair of triplets (*s*_1_, *p*_1_, *o*_1_, *s*_2_, *p*_2_, *o*_2_) indicating the two ends of the breakpoint:

1. chromosomes in the reference genome (*s*_1_, *s*_2_) (both *s*_1_, *s*_2_ *∈* ℛ𝒢);
2. starting or ending positions of each interval (*p*_1_, *p*_2_); and
3. and orientations (*o*_1_, *o*_2_) (both *o*_1_, *o*_2_ *∈ {*+, −*}*)

The positions *p*_1_ and *p*_2_ denote starting or ending positions, depending on the orientation. Specifically:

1. if *o*_1_ = + and *o*_2_ = +, b describes a junction between genomic intervals *s*_1_[…, *p*_1_] and *s*_2_[…, *p*_2_].
2. if *o*_1_ = + and *o*_2_ = −, b describes a junction between genomic intervals *s*_1_[…, *p*_1_] and *s*_2_[*p*_2_, …].
3. if *o*_1_ = − and *o*_2_ = +, b describes a junction between genomic intervals *s*_1_[*p*_1_, …] and *s*_2_[…, *p*_2_].
4. if *o*_1_ = − and *o*_2_ = −, b describes a junction between genomic intervals *s*_1_[*p*_1_, …] and *s*_2_[*p*_2_, …].

We note that (*s*_1_, *p*_1_, *o*_1_, *s*_2_, *p*_2_, *o*_2_) and (*s*_2_, *p*_2_, *o*_2_, *s*_1_, *p*_1_, *o*_1_) describe the same breakpoint. To facilitate comparisons over breakpoints, we therefore require either *s*_1_ ≺ *s*_2_ or *s*_1_ = *s*_2_, *p*_1_ < *p*_2_. It is also possible, though rare, that a breakpoint connects (*s*_1_, *p*_1_, +) to (*s*_1_, *p*_1_, −), representing a single nucleotide duplication. As such we ignore these cases.

#### Breakpoint Graph

A **Breakpoint Graph** is a weighted, undirected graph 𝒢 = (*V, E* = *E*_*s*_ ∪ *E*_*c*_ ∪ *E*_*d*_, CN) that contains information pertaining to breakpoints in a sample. We define the entities in this graph as follows:

*V* (*V* ⊆ ℛ *𝒢 ×* ℕ *× {*+, −*}*) represents either the starting or ending position of a genomic interval (except the two special source nodes *s* and *t*, as described below);

*E*_*s*_ represents **sequence edges**, or connections between the starting and ending positions in a genomic interval;

*E*_*c*_ represents the **concordant edges** that connect two consecutive genomic intervals *s*_*i*_[…, *p*] and *s*_*i*_[*p* + 1, …].

*E*_*d*_ represents the **discordant edges** connecting two nodes *v*_1_ = (*s*_1_, *p*_1_, *o*_1_) and *v*_2_ = (*s*_2_, *p*_2_, *o*_2_) where *s*_1_*≠ s*_2_, or |*p*_1_ − *p*_2_| > 1, or *o*_1_ = *o*_2_. A discordant edge could also connect a node *v* = (*s*_1_, *p*_1_, *o*_1_) to itself - we call such discordant edges **foldbacks**. Foldback edges will introduce self-loops in a breakpoint graph. We sometimes refer to a **breakpoint edge** as either a concordant or discordant edge in the breakpoint graph, indicating a discontinuity on the reference genome.

CN CN maps each edge to a fractional **copy number**: CN : *E* → ℚ+.

Note that in a breakpoint graph, each node is connected to a single sequence edge, and a single concordant edge as well; but it may connect to multiple discordant edges. Similar to (Deshpande et al. 2019; Hadi et al. 2020; Aganezov and Raphael 2020), we require that the CN assignment is “balanced” for each sequence edge (*u, v*) *∈ E*_*s*_, i.e., the sum of CN values from the breakpoint edges connected to (*u, v*) equals to the CN of thos sequence edge. A node connected to a foldback edge receives two times the CN from the foldback edge it connects to when summing up CN values from discordant edges.

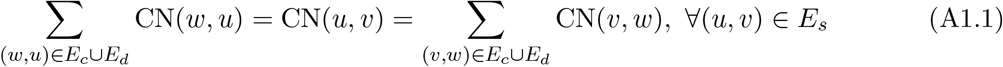

As per Amplicon Architect (AA) (Deshpande et al. 2019), **amplified intervals** (or **amplicon intervals**) describes a set ℐ = *{*(*s*_*i*_, *l*_*i*_, *r*_*i*_)*}* as the union of genome intervals represented by all sequence edges in a breakpoint graph - where *l*_*i*_ gives the smallest coordinate among a few sequence edges connected by concordant edges; and *r*_*i*_ gives the largest coordinate among a few sequence edges connected by concordant edges. Each string *s*_*i*_ in ℛ𝒢 can include multiple, non-adjacent amplified intervals: for 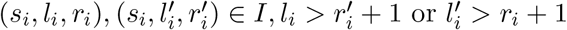. The simplest non-empty breakpoint graph gives a single amplified interval, a single sequence edge, and no breakpoint (concordant/discordant) edges. We refer to an **amplicon** as the union of all amplified intervals and their (discordant) connections. In other words, an amplicon corresponds to a breakpoint graph. In a tumor sample there could be multiple amplicons whose intervals are non-intersecting.

In addition to the nodes representing the starting or ending position of a genomic interval (sequence edge), we introduce two artificial nodes, *s ∈ V* and *t ∈ V* in each breakpoint graph 𝒢. They both connect to the end of each amplified interval, as well as a sequence edge which is only connected to another sequence edge with smaller CN by concordant edges, and therefore is deemed to violate the balanced CN constraint A1.1 without the source connections. Edges connected to source nodes are treated as discordant edges.

A **walk** in a breakpoint graph *𝒢* is a sequence of nodes *v*_1_, *v*_2_, …, *v*_*w*_, where for all 1 ≤ *i* < *w*, (*v*_*i*_, *v*_*i*+1_) *∈ E*, and the edges alternate between sequence and breakpoint edges (i.e. if (*v*_*i*_, *v*_*i*+1_) *∈ E*_*s*_ then (*v*_*i*−1_, *v*_*i*_) *∈ E*_*c*_ ∪ *E*_*d*_ and (*v*_*i*+1_, *v*_*i*+2_) *∈ E*_*c*_ ∪ *E*_*d*_). A *path* is a walk with no node repeated (*v*_*i*_ = *v*_*j*_ ⇔ *i* = *j*). An *s, t***-walk/path** in a walk or path starts at *s* and ends at *t*. A **cycle** or **cyclic walk** is a walk where the first edge starts with and the last edge ends with the same node, i.e., *v*_1_ = *v*_*w*_*≠ s, t*. The cycle is **simple** if no node except the first/last one is repeated.

With these definitions, we say that a properly reconstructed breakpoint graph 𝒢 represents a superimposition of all ecDNAs and rearranged genomes. These rearranged genomes are represented by walks of alternating sequence edges and breakpoint edges. For example, an *s, t*-path of alternating sequence and breakpoint edges may represent a linear focal amplification; ecDNA form cycles of alternating sequence and breakpoint edges.

### A2. Breakpoint Graph Reconstruction from Long Reads

#### Seed Interval Detection

CoRAL detects seed amplified intervals from whole genome CNV calls of mapped long reads (e.g., with third party tools like CNVkit (Talevich et al. 2016)). In fact, CNV calls give a partition of the reference genome ℛ𝒢 into non-overlapping genomic intervals, each with a distinct copy number. We first remove centromeric intervals and select candidate seed intervals as the intervals *a*_*i*_ whose copy number is at least max(4.0 + `CN_chr_arm` (*a*_*i*_), 6.0), where `CN_chr_arm` (*a*_*i*_) is the average (length-weighted) copy number of all intervals on the same chromosome arm as *a*_*i*_. We merge adjacent candidate intervals and candidate intervals on the same chromosome and within 2 * 10^5^bp (= 2*δ*, see below) distance. Among the merged intervals we finally select the ones with aggregated size at least 10^5^bp (= *δ*) as seed intervals. If there are no seed amplified intervals, CoRAL stops by reporting no focal amplifications in the input sample. Otherwise CoRAL proceeds with amplified interval search, breakpoint graph construction and cycle decomposition.

#### Amplified Interval Search

Let Δ be the maximum length of (amplified) genomic segment in an amplicon without breakpoints in the middle. According to TCGA and PCAWG data from (Kim et al. 2020), we set Δ to 10^6^bp. Let *δ* be the size of flanking region surrounding an amplified interval. Following from (Deshpande et al. 2019) we set *δ* to 10^5^bp. We present the pseudocode of amplified interval search, given a seed interval list ℐ_*s*_, as follows.

#### Breakpoint Clustering

Define a “match” between two breakpoints bp_1_ = (*s*_1_, *p*_1_, *o*_1_, *s*_2_, *p*_2_, *o*_2_) and 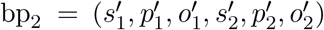 when they have the same chromosome, orientation, and close positions, i.e., 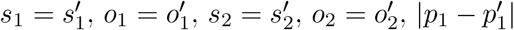 ≤ `bp_distance_cutoff` and 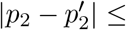 `bp_distance_cutoff`, for some `bp_distance_cutoff`. We cluster a collection of breakpoints *B* through two steps. First we compute crude clusters with the following greedy strategy and a large `bp_distance_cutoff` = 2000bp (the largest possible distance given by the chimeric alignments between two reads which support a single breakpoint - see (Sedlazeck et al. 2018)): start with an empty set of clusters; for each breakpoint bp, if there exists a cluster 𝒞 containing another breakpoint bp’ which matches bp, then add the breakpoint bp to cluster 𝒞; otherwise start a new cluste 𝒞^*′*^ and add bp to 𝒞^*′*^.

We then refine each crude cluster and compute the exact breakpoint. For each cluster 𝒞 resulting from the above step, we compute the average *μ*_1_ and standard deviation *σ*_1_ of the first (“smaller”) position *p*_1_, as well as *μ*_2_ and *σ*_2_ of the second (“larger”) position *p*_2_. Then we remove the ‘outlier’ breakpoints in 𝒞 with a smaller `bp_distance_cutoff` = 100bp, when *p*_1_ < *μ*_1_ − max(3 * *σ*_1_,`bp_distance_cutoff`), or *p*_1_ > *μ*_1_ + max(3 * *σ*_1_,`bp_distance_cutoff`), or *p*_2_ < *μ*_2_ − max(3 * *σ*_2_,`bp_distance_cutoff`), or *p*_2_ > *μ*_2_ + max(3 * *σ*_2_,`bp_distance_cutoff`). If the remaining cluster size is still at least `bp_clustersize_cutoff`, we keep the breakpoint corresponding to cluster 𝒞, otherwise we split 𝒞 into four subclusters: (i) breakpoints with *p*_1_ < *μ*_1_ and *p*_2_ < *μ*_2_; (ii) *p*_1_ < *μ*_1_ and *p*_2_ > *μ*_2_; (iii) *p*_1_ > *μ*_1_ and *p*_2_ < *μ*_2_; and (iv) *p*_1_ > *μ*_1_ and *p*_2_ > *μ*_2_. The subclusters with size at least `bp_clustersize_cutoff`are repeated with the above procedure and otherwise are discarded. The final breakpoint positions for a cluster 𝒞 is determined by the *mode* of *p*_1_ and *p*_2_ in 𝒞. If there are multiple modes then we use the average positions. The `bp_clustersize_cutoff` is determined by the maximum of 3 and haploid coverage to avoid false positive chimerisms in long read alignments. See below how CoRAL estimates the haploid coverage in a tumor sample.

##### Algorithm 1 AmplifiedIntervalSearch(ℐ_*s*_, *δ*, Δ, bp_clustersize_cutoff) ▹ ℐ_*s*_: seed intervals

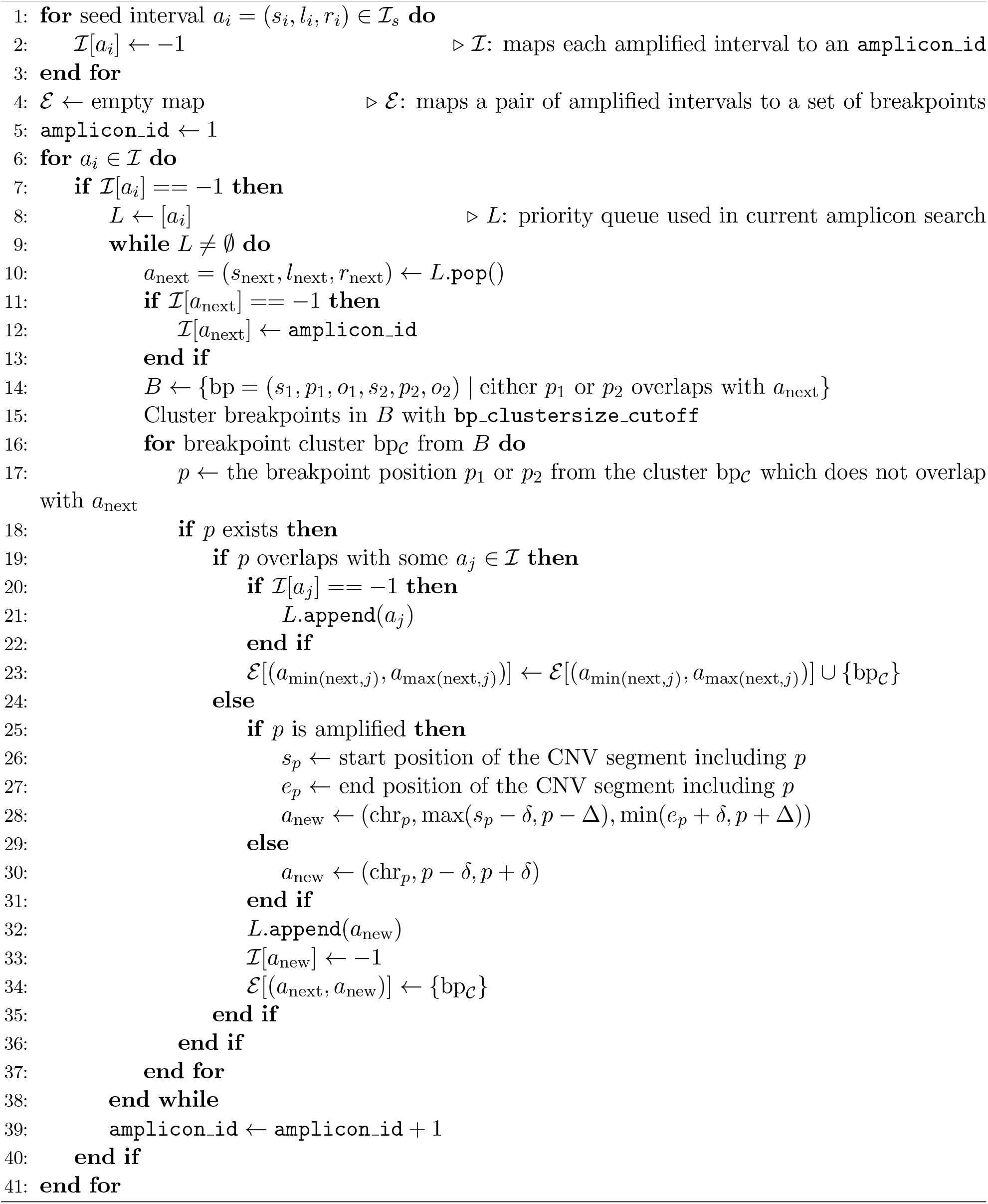

### A3. Copy Number Assignment

#### Estimating diploid coverage

The CN assignment of CoRAL requires an estimation of diploid coverage *θ*_LR_. The estimation can be derived from CNV calls used for detecting seed amplified intervals, with the assumption that majority of the donor genome is not amplified. Recall that CNV calls give a partition of the reference genome ℛ𝒢 into non-overlapping genomic intervals, each with a distinct copy number. To estimate *θ*_LR_ CoRAL first sorts these intervals according to their predicted copy numbers, and locate the interval *a*_*i*_ = (*s*_*i*_, *l*_*i*_, *r*_*i*_) at the 40-th percentile in the sorted list. If the length of *a*_*i*_ is less than 10^7^bp, CoRAL iteratively includes the interval *a*_*i*−1_ and *a*_*i*+1_ in the sorted list, until the aggregate size of the included intervals is at least 10^7^bp. CoRAL computes *θ*_LR_ as the (length-weighted) average long read coverage of all selected intervals centered at *a*_*i*_.

#### Maximum likelihood CN assignment

Given *θ*_LR_, we model the total number of nucleotides *N*_*e*_ on each sequence edge *e ∈ E*_*s*_ as a normal distribution with mean and variance both *θ*_LR_ · CN(*e*) · *l*(*e*), where *l*(*e*) denotes the length (in bp) of the sequence edge

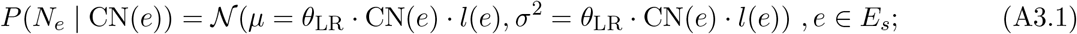

and the number of reads 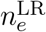 supporting each concordant and discordant edge *e ∈ E*_*c*_ ∪ *E*_*d*_ as a Poisson (similar to (Medvedev et al. 2010; Deshpande et al. 2019)) with mean *θ*_LR_ · CN(*e*)

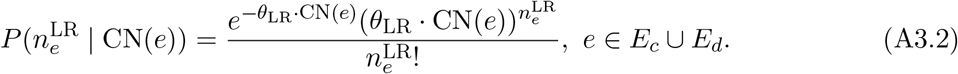

To estimate CN, CoRAL computes the maximum likelihood ℒ of CN using the joint distribution of observed number of nucleotides on each sequence edge and the observed read counts on each concordant/discordant edge

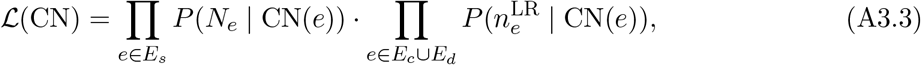

with the constraint that CN is balanced for each node *v* (by rewriting equation A1.1), i.e.,

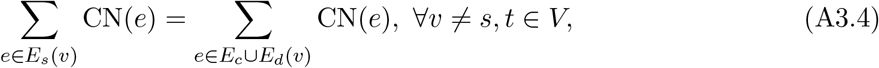

where *E*_*s*_(*v*), *E*_*c*_(*v*), and *E*_*d*_(*v*) stand for the sequence edge, concordant edge, and discordant edges connected to node *v*, respectively. Edges connected to the source nodes *s* and *t* do not contribute to the likelihood (objective) function, nor to the constraints. The (convex) optimization problem was solved using CVXOPT package (https://github.com/cvxopt/cvxopt).

#### A4. Cycle Decomposition

In cycle decomposition we decompose an amplicon 𝒢 into a collection of cycles and *s, t*-walks, with high copy numbers. For all sequence edges (*u, v*) *∈ E*_*s*_, define the *length-weighted-copy-number* using C_*l*_(*u, v*) = CN(*u, v*) · *l*(*u, v*), where *l*(*u, v*) denotes the length (in bp) of the corresponding segment. Similarly, for graph 𝒢.

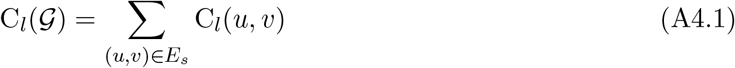

The MIQCP for cycle extraction works with 3 parameters: *k* as the maximum number of walks; *α* as the minimum fraction of length-weighted copy number explained, and *β* as the minimum fraction of path constraints satisfied. Of these, *k* is learned starting with *k* = 1, according to two modes. In the **FullQP** mode, the MIQCP attempts a solution with at most *k* walks that satisfy other constraints, or returns ‘infeasible.’ The value of *k* is doubled until feasibility is reached or *k* > |*E*|. The **greedy mode** is described below. We implement both quadratic programs with through the python3 interface of Gurobi 10.0.1.

We use the following **key** variables.

- *w*_*i*_ *∈* ℚ ≥ 0: denotes the copy number for walk *W*_*i*_ (1 ≤ *i* ≤ *k*); an auxiliary variable *z*_*i*_ *∈ {*0, 1*}* indicates if *w*_*i*_ > 0;
- *x*_*uvi*_ *∈* ℤ ≥ 0 represents the number of times walk *W*_*i*_ traverses (*u, v*) for each edge (*u, v*) *∈ E* and 1 ≤ *i* ≤ *k*;
- *P*_*j*_ *∈ {*0, 1*}* indicates if subwalk constraint *p*_*j*_ is satisfied for 1 ≤ *j* ≤ *m* ;

The MIQIP(*k, α, β*) objective is given by:

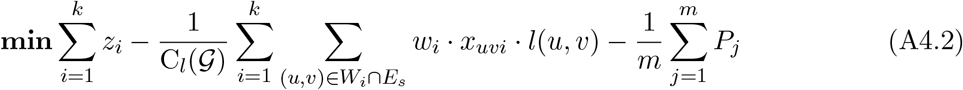

subject to the constraints:

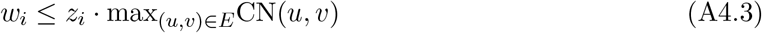

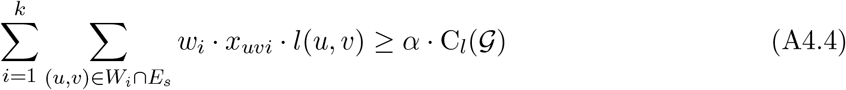

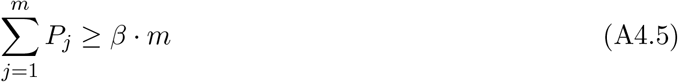

In addition, the MIQIP satisfies a number of auxiliary constraints that constrain the cycles and walks to satisfy normal definitions, and those require the following auxiliary variables:

- *c*_*vi*_ *∈ {*0, 1*}* for node *v ∈ V* − *{s, t}* and 1 ≤ *i* ≤ *k*. *c*_*v,i*_ = 1 iff the walk *W*_*i*_ forms a cycle starts (and ends) with node *v*, if *W*_*i*_ exists;
- *p*_*ij*_ *∈ {*0, 1*}*, for 1 ≤ *i* ≤ *k*, 1 ≤ *j* ≤ *m* indicates if subwalk constraint *p*_*j*_ is satisfied by walk *W*_*i*_;
- 0 ≤ *d*_*vi*_ ≤ |*V* | for each node *v ∈ V*, 1 ≤ *i* ≤ *k*, starting with a number 1 for the initial node and incrementing for the next node in the cycle/walk. It is used to ensure connectivity;
- 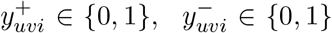 for each edge (*u, v*) *∈ E* and 1 ≤ *i* ≤ *k*, is also used to ensure connectivity.

#### Additional Constraints

1. Each *W*_*i*_ should form a valid walk of alternating sequence and breakpoint edges. In other words, for each node *v ∈ V* −*{s, t}*, the sum of *x*_*uvi*_ from sequence edges (*u, v*) *∈ E*_*s*_ it connects to should equal to the sum of *x*_*uvi*_ from breakpoint edges (*v, w*) *∈ E*_*c*_ ∪ *E*_*d*_ it connects to.

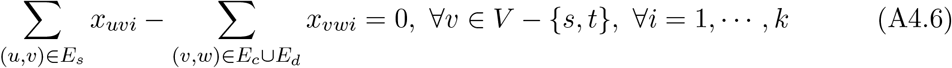
2. The total CN of all cycles/walks passing through an edge (*u, v*) *∈ E* is at most CN(*u, v*).

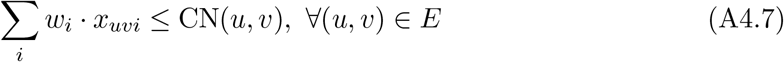
3. We require that each cycle/walk traverses through a discordant edge (*u, v*) *∈ E*_*d*_ at most *R*(*u, v*) times, otherwise cycles were not possible.

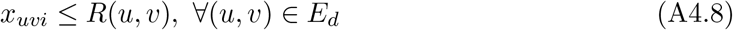

*R*(*u, v*) can be a small, fixed number for all (*u, v*) *∈ E*_*d*_, e.g., 2 to allow a discordant edge to be traversed in each direction as the graph is undirected. However, by default CoRAL computes an empirical *R*(*u, v*) for each (*u, v*) *∈ E*_*d*_ by clustering the number of reads 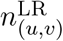 supporting the discordant edge with the constraint that two observations 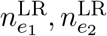 must belong to different clusters if one of them is at leats 5 times larger than the other, i.e., 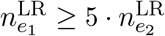 or 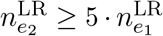. In case there exists some edges with *R*(*u, v*) > 1, the constraint A4.11 below prevents every edge in a cycle being repeated multiple times.
4. Each walk *W*_*i*_ either forms a cycle starting at node *v*, or starts at *s* and ends at *t* if it exists. If *W*_*i*_ forms a cycle we require that there exists one concordant or discordant edge connected to *c*_*vi*_ which occurs only once in the cycle.

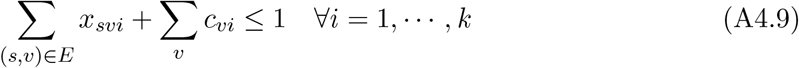

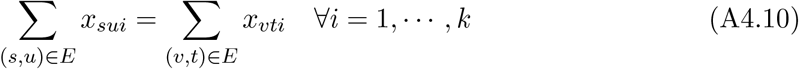

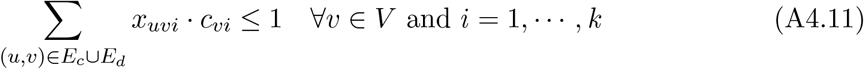
5. *x*_*uvi*_ and *z*_*i*_ are consistent. In other words *z*_*i*_ = 1 if and only if there exists some *x*_*uvi*_ > 0. Since *x*_*uvi*_ are not binary, the consistency between *x*_*uvi*_ and *z*_*i*_ can be guaranteed through 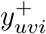 or 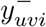

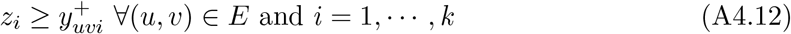

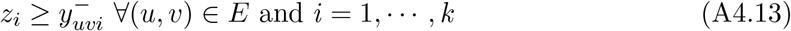

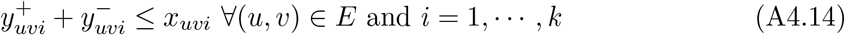
6. Connectivity. The idea is to use *d*_*vi*_ to encode the “discovery order” of the nodes in walk *W*_*i*_. If *W*_*i*_ is a cycle then we start with the node *v* where *c*_*vi*_ = 1; otherwise we start with the source node *s*. *d*_*vi*_ for the starting node *v* is set to 1. Each node *v* (except the starting node) in *W*_*i*_ is assumed to be “discovered” **uniquely** by another already “discovered” node *u* through edge (*u, v*), satisfying *d*_*vi*_ ≥ *d*_*ui*_ + 1. As breakpoint graph is undirected, we assume an order < of nodes and for each edge (*u, v*) we introduce two binary variables 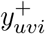 or 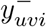 indicating respectively the larger node in *u, v* is discovered from the smaller node through edge (*u, v*); or the smaller node is discovered from the larger node through edge (*u, v*). Specifically, if *v* is discovered from *u* and *v* > *u* or *u* is discovered from *v* and *v* < *u* then 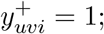 if *v* is discovered from *u* and *v* < *u* or *u* is discovered from *v* and *v* > *u* then 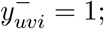 in all other cases 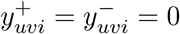 . The nodes do not belong to *W*_*i*_ also have *d*_*vi*_ = 0 and 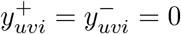 We first set up the constraint for the starting node.

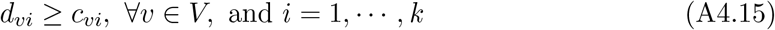

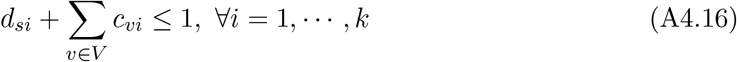

We require that *d*_*vi*_ = 0 if node *v* is not a part of *W*_*i*_.

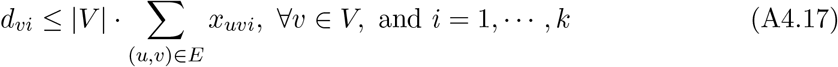

Fold-back edges (self-loops) can not have positive 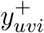 or 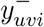.

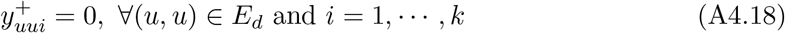

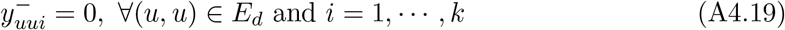

Each node can be discovered from at most one neighboring node.

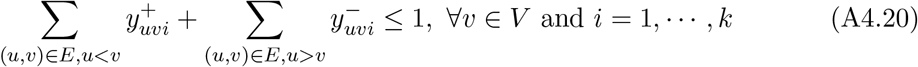

If a node *v* is included in walk *W*_*i*_ and it is not the starting node of a cycle, then it must be discovered through some 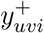 or 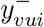.

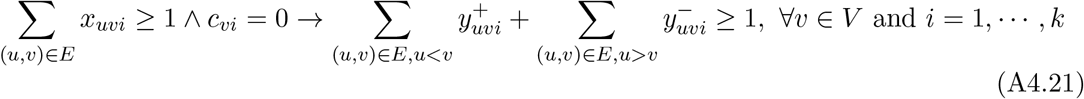

Equation (A4.21) can be rewritten as a quadratic constraint, as follows. For all *v ∈ V* and *i* = 1, …, *k*,

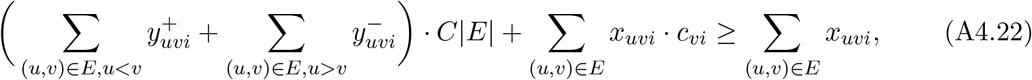

where *C* is a large constant representing the maximum number of times that any edge (*u, v*) *∈ E* can be traversed by a cycle or walk (e.g., *C* can be set to max degree of 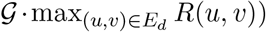. Finally, we connect 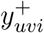 and 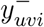 with *d*_*vi*_. If a node *v* is included in walk *W*_*i*_ and it is not the starting node of a cycle, then *d*_*vi*_ ≥ *d*_*ui*_ + 1 for the node *u* it was discovered from. For all *v ∈ V* and *i* = 1, …, *k*

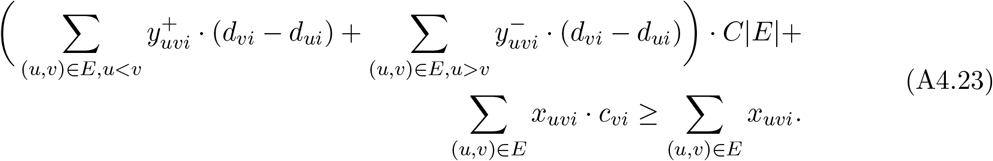
7. **Subwalk constraints**. We enforce a weak constraint by requiring each walk *p*_*j*_ *∈* 𝒫 as a subgraph of the graph induced by some walk *W*_*i*_.

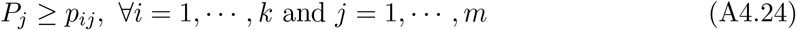

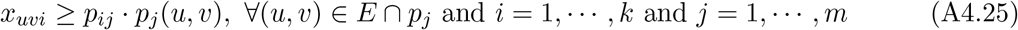

where *p*_*j*_(*u, v*) the number of times walk *p*_*j*_ passes through an edge (*u, v*).

#### Subwalk constraints from long reads

To compute subwalk constraints we extract all long reads mapped within the amplified intervals defined by the breakpoint graph 𝒢, and map each of them to 𝒢. We filter out reads which could not be fully mapped to the breakpoint graph, due to additional breakpoints or partially non-overlap with any sequence edge. Each of the remaining reads mapped to 𝒢 should give a walk in 𝒢 starting and ending with sequence edges. We further filter out walks where the first or last sequence edge overlaps with the corresponding read by less than 500bp; and then walks with at most 3 edges (which only cover one breakpoint). Finally, we filter out walks that form a subwalk of any other walk and return the remaining walks as the subwalk constraints 𝒫 to be used in cycle extraction, either full QP or greedy QP.

#### MIQIP-greedy

Let 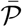= *{j* | path *p*_*j*_ is not satisfied by any previously selected walk*}*. The full greedy MIQCP to identify the next walk *W*_*i*_ is given by:

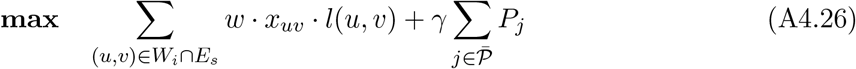

subject to

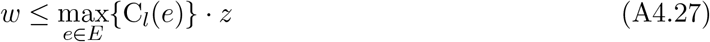

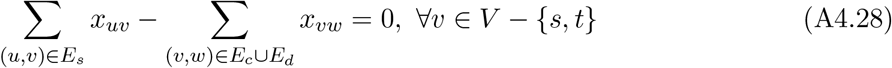

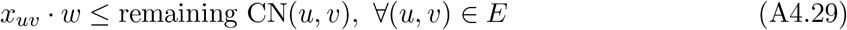

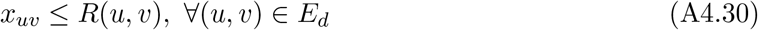

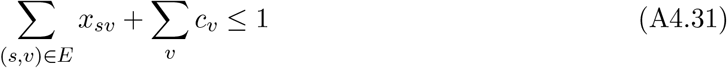

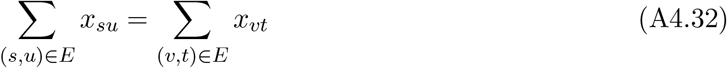

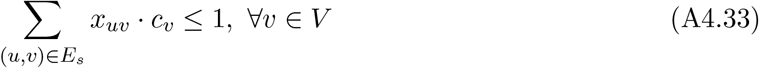

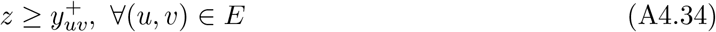

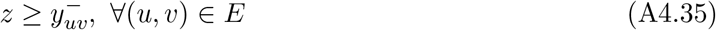

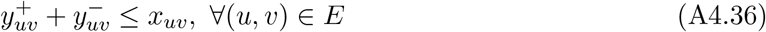

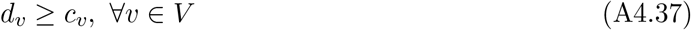

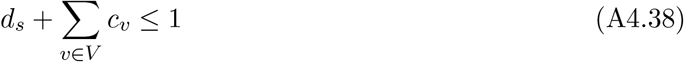

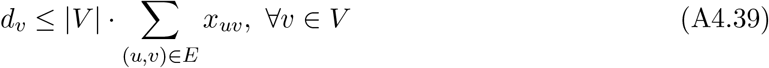

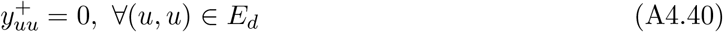

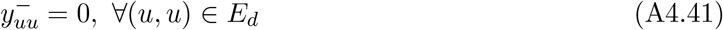

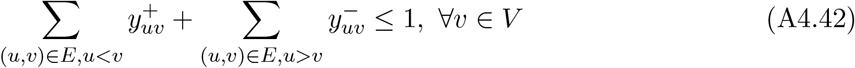

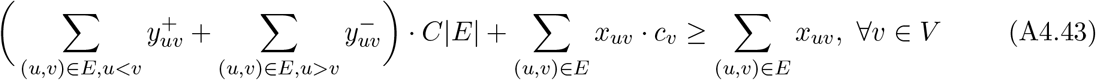

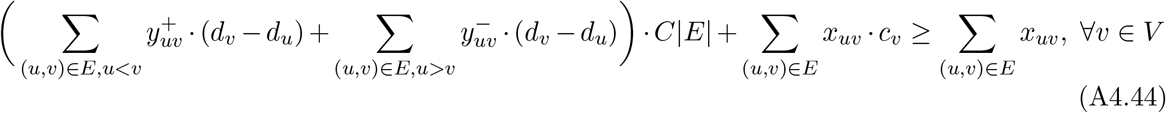

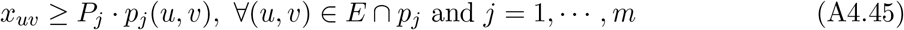

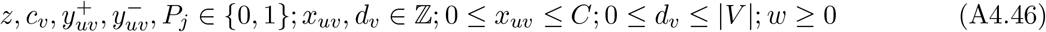

### A5. Simulating amplicon structures with ecSimulator

We utilized an updated version of ecSimulator (Luebeck et al. 2020) (version 0.6.0, https://github.com/AmpliconSuite/ecSimulator) to simulate ecDNA genome structures derived from three different contexts and simulated both long and short reads from those structures at varying copy number levels.

In brief, ecSimulator uses a user-specified YAML input to set simulation parameters, allowing the user to specify properties such as the number of genomic intervals in the simulated ecDNA, the number of possible locations of breakpoints, and the rates of SV types (deletion, duplication, inversion, translocation, foldback). We utilized ecSimulator’s default parameters for SV type frequency. ecSimulator supports the simulation of ecDNA derived from three different modes of genesis. First, the episome model, whereby the structure is initialized with only head-to-tail closure of the interval(s). Second, the two-foldback model where the interval is bound on left and right by a foldback SV (inverted duplication). Third, a chromothriptic model, whereby intervals are separated by a deletions and then closed head-to-tail, simulating the oscillating CN states observed in chromothripsis. To simulate the internal rearrangements of the genome structure observed on ecDNA intervals, ecSimulator first assigns *n* breakpoints randomly throughout the intervals according to the number of possible breakpoints desired by the user. By pre-assigning possible breakpoint locations, it is possible to easily create structures containing multiple copies of a breakpoint or to enable breakpoint re-use. ecSimulator then performs multiple rounds of SV boundary assignment and rearrangement, with the default number of rounds being 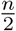 . A random number of consecutive sub-segments of the intervals are selected, where sub-segments are defined as pieces of the intervals separated by the pre-assigned breakpoint locations. The selected sub-segments then undergo random assignment of the SV type being applied, and the assigned SV type is simulated inside the structure on those selected segments.

To simulate reads we maintained the following default parameters:

Target_size: 2,000,000

Mean_segment_size: 150,000

num_intervals: “auto”

same_chromosome: False (allowing segments to be recombined across chromosomes)

allow_interval_reuse: True (allowing higher multiplicity in an amplicon)

viral_insertion: False

del: 0.6 (probability of deletion)

dup: 0.5 (probability of duplication)

inv: 0.4 (probability of inversion)

trans: 0.4 (probability of translocation)

fback: 0.05 (probability of an inverted duplication)

In addition to these default parameters, we simulated amplicons from the chromothripsis, episomal, or two-foldback origins. For each origin, we simulated 5 replicates of an amplicon with 1, 3, 5, 10, or 20 breakpoints resulting in a dataset of 75 simulated amplicons.

### A6. Whole Genome Sequencing (WGS)

#### Public data sources

We obtained sequencing data from the following public repositories. Illumina data for COLO320-DM (Wu 2019) was obtained from SRX5055021; Illumina data for COLO320-HSR (Wu 2019) was obtained from SRX5930165. Illumina data for GBM39 (Wu 2019) was obtained from SRX5055022; Illumina data for GBM39-HSR (Wu 2019) was obtained from SRX5930166; Illumina data for CHP-212 was obtained from SRX8044100; Illumina data for COLO320-DM (Hung 2021) was obtained from SRX11096731. Long-read Nanopore data for COLO320-DM (Hung 2021) was obtained from SRX9346575; Long-read Nanopore data for CHP-212 was obtained from SRX8044102.

We additionally sequenced the other 4 Illumina samples (COLO320-DM (mono), GBM39 (mono), PC3-DM (mono), PC3-HSR (mono)) and 8 Nanopore samples (COLO320-DM (Wu 2019), COLO320-HSR (Wu 2019), GBM39 (Wu 2019), GBM39-HSR (Wu 2019), COLO320-DM (mono), GBM39 (mono), PC3-DM (mono), PC3-HSR (mono)), as described in the following sections (see “Short-read WGS” and “Nanopore Long-read WGS”).

#### Deriving monoclonal cell lines

The following approaches were taken to derive stable monoclonal cell lines for this study:

1. **PC3:** PC3 cell line with a mixed pooled of cells containing *MYC* amplification as either ecDNA (for PC3-DM) or HSR (for PC3-HSR) were seeded onto a 96-well plate at a density of 0.5 cell/well, in an attempt to have a maximum of one cell seeded per well. Cells were then expanded and iteratively expanded into a 24-well plate, 6-well plate, then 10 cm dish over the course of one month. PC3 cells were maintained in Dulbecco’s modified Eagle’s medium (DMEM; Corning, #10-013-CV) supplemented with 10% fetal bovine serum (FBS; Hyclone, SH30396.03) and 1% pen-strep (PS; Thermo Fisher Scientific, 15140-122).
2. **GBM39:** GBM39 cell line with a mixed pooled of cells containing *EGFR*vIII amplification as ecDNA were sent to Cell Microsystems for monoclonal expansion. In brief, cells were subjected to TrypLE digestion to yield single cells, and each single cell was seeded onto a CellRaft Array using the CellRaft AIR® System. The cells were maintained and expanded as a monoclonal line. Neurosphere culture medium Dulbecco’s modified Eagle’s medium/nutrient mixture F-12 (DMEM/F12 1:1; Gibco, 11320-082) with 1% PS, GlutaMAX (Gibco, 35050061), B27 supplement (Gibco, 17504044), 20 ng/ml epidermal growth factor (EGF; Sigma-Aldrich, E9644), 20 ng/ml fibroblast growth factor (FGF; Peprotech) and 5 *μ*g/ml (Sigma-Aldrich, H3149-500KU) was used throughout to maintain the culture.
3. **COLO320-DM:** A recently monoclonalized cell line of COLO320-DM was received as a gift from the Mischel group.

#### Short-read WGS

WGS libraries were prepared by DNA tagmentation. We first transposed it with Tn5 transposase produced as previously described, in a 50-*μ*l reaction with TD buffer, 10ng DNA and 1 *μ*l transposase. The reaction was performed at 50°C for 5 minutes, and transposed DNA was purified using Zymo DNA Clean & Concentrate kit (Zymo, 1159U33). Libraries were generated by 7 rounds of PCR amplification using NEBNext High-Fidelity 2× PCR Master Mix (NEB, M0541L), purified using SPRIselect reagent kit (Beckman 635 Coulter, B23317) with double size selection (0.8× right, 1.2× left) and sequenced on the Illumina Nextseq 550 platform. Reads were trimmed of adapter content with Trimmomatic (Bolger et al. 2014) (version 0.39), aligned to the hg38 genome using BWA MEM (Li and Durbin 2009) (version 0.7.17-r1188), and PCR duplicates removed using Picard’s MarkDuplicates (version 2.25.3).

#### Nanopore Long-read WGS

We performed default long-read sequencing on COLO320-HSR, GBM39 (mono), and PC3-DM (mono). To do so, high molecular weight (HMW) genomic DNA from approximately 2 million cells was extracted using the Qiagen Puregene DNA Kit (Qiagen 158023) and prepared for long-read sequencing using the Oxford Nanopore Ligation Sequencing Kit V14 (Oxford Nanopore Technologies SQK-LSK114) according to the manufacturer’s instructions.

We performed Ultralong seqeuncing on COLO320-DM (mono) and PC3-HSR (mono). To do so, HMW genomic DNA was extracted from approximately 6 million cells using the NEB HMW DNA Extraction Kit for Cells & Blood (NEB T3050) and prepared for sequencing using the Oxford Nanopore Ultra-Long DNA Sequencing Kit V14 (Oxford Nanopore Technologies SQK-ULK114) according to the manufacturer’s instructions. Libararies were sequenced on a PromethION (Oxford Nanopore Technologies) using a 10.4.1 flow cell (Oxford Nanopore Technologies FLO-PRO114M). Basecalling from raw POD5 files was performed using Dorado (Oxford Nanopore Technologies, version 0.2.1) and aligned to GRCh38 using minimap2 (Li 2018).

In addition, we performed long-read sequencing on non-monoclonalized GBM39 and GBM39-HSR as follows: HMW genomic DNA was extracted using a MagAttract HMW DNA Kit (Qiagen 67563) and prepared for long-read sequencing using a Ligation Sequencing Kit (Oxford Nanopore Technologies SQK-LSK109) according to the manufacturer’s instructions. Libraries were sequenced on a PromethION (Oxford Nanopore Technologies) using a 9.4.1 flow cell (Oxford Nanopore Technologies FLO-PRO002). Bases were called from FAST5 files using Guppy (Oxford Nanopore Technologies, v.2.3.7) and read were aligned to GRCh38 using minimap2 (Li 2018).

### A7. Amplicon comparison statistics

#### Breakpoint-graph accuracy

We evaluate the proportion of breakpoint edges captured for a reconstruction as follows: let 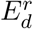 be the set of discordant edges in a reconstructed breakpoint graph 𝒢_*r*_ and 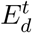 be the set of discordant edges in a simulated breakpoint graph *𝒢*_*t*_. For each edge 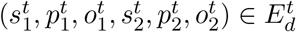 we evaluate if a similar edge appears in the set of reconstructed edges 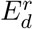. Specifically, an an edge is deemed similar if the orientations are identical and the positions differ by at most 100bp - i.e., 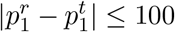 and 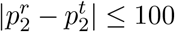, where 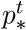 is the true coordinate and 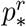 is the reconstructed coordinate. The proportion of true edges recovered is reported.

#### Cycle-interval overlap

Let 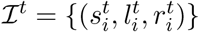 be the set of intervals covered by the simulated cycle and let 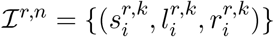 be the set of intervals covered by the *n*-th cycle returned by a reconstruction algorithm. (Recall that CoRAL and AA return a set of possible cycles). Let *δ*(*ℐ*_*i*_, *ℐ*_*j*_) be a function that returns the length (in nucleotides) of the overlapping region between two intervals *ℐ*_*i*_ and *ℐ*_*j*_ (if *s*_*i*_*≠ s*_*j*_ then *δ*(*ℐ*_*i*_, *ℐ*_*j*_) = 0); let *ℓ*(*ℐ*_*j*_) indicate the length (in nucleotides) of the interval *ℐ*_*j*_ (i.e., |*r*_*i*_ − *l*_*i*_|). Then the Cycle-interval overlap is for the *n*-th cycle is defined as

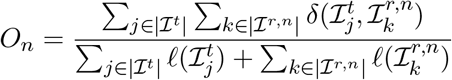

For a given simulated amplicon, we report best overlap statistic found across all returned reconstructions.

#### Cyclic Longest Common Subsequence (LCS)

Let *W*_*t*_ be a true, simulated cycle defined by an <u>ordered</u> sequence of intervals 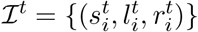, and let 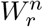 be the *n*-th cycle defined by the <u>ordered</u> sequence of intervals 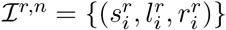 returned by a cycle decomposition algorithm (as above, recall that CoRAL and AA return a set of possible cycles). Let LCS(*I*^*t*^, *I*^*r,n*^(*j*)) be defined as the longest common subsequence between the two ordered interval sets (in nucleotides). Note that the common subsequence does not need to be contiguous. For example, for the sequence *ℐ*_1_ = (1, 2, 3, 4, 5) and *ℐ*_2_ = (1, 3, 4, 5, 6), the LCS would be (1, 3, 4, 5).

To account for cycle rotation, we also consider the rotated cycle *W* (*j*) containing *m* intervals where the interval set is rotated around the index *j*: ℐ = *{*ℐ_*j*_, ℐ_*j*+1_ … ℐ_*m*_, ℐ_1_, … ℐ_*j*−1_*}*. In addition, we consider the reverse cycle whereby 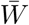 consists of the reverse of all intervals: 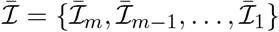 and 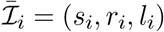.

Then, the cyclic longest common subsequence for a reconstructed cycle consisting of *m* intervals is defined as (only considering intervals that overlap between the true and reconstructed cycle):

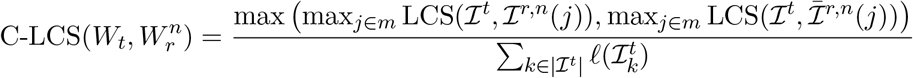

In words, this measure reports the longest common subsequence that can be found between the two cycles *W*_*t*_ and 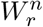 while considering all possible rotations and reversals, and normalized to the length of the true cycle (in nucleotides).

#### Reconstruction Length Error

This statistic measures the differences in reconstructed cycle length and true cycle length. As before, let *W*_*t*_ be a true, simulated cycle defined by an <u>ordered</u> sequence of intervals 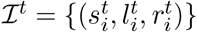, and let 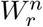 be the *n*-th cycle defined by the <u>ordered</u> sequence of intervals 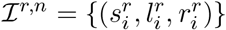. Furthermore, let *L*_*_ be the length of cycle *W*_*_ (where * indicates *t* or some reconstruction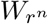 )): 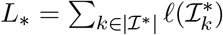

Then, the reconstruction length error for a particuler reconstruction 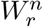 is defined as

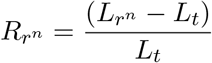

The final value *R*_*r*_ is defined as the value for the best reconstruction: 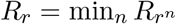. For clarity of presentation, we report log_2_(1 + *R*_*r*_).

#### *K*−Heaviest Cycle Weight Ratio

This statistic is agnostic to the true cycle and reflects the “entropy” of a given reconstruction, by measuring the fraction of total copy-number can be accounted for by the *k* heaviest cycles. Here, the length-weighted copy-number of the entire breakpoint graph (denoted as C_*l*_(𝒢)) is calculated from the sequence edges *E*_*s*_ reported in a breakpoint graph 𝒢 and is defined in Eqn.(2) of the main text. The length-weighted copy-number of a particular cycle is defined as the sum of the length-weighted copy-number the intervals reported in a particular reconstructed cycle 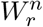 that is defined by *m* total intervals contained in the set ℐ ^*r,n*^. Using similar notation as above, if C_*l*_(*u, v*) denotes the length-weighted copy-number of a particular interval covering (*u, v*), then we define the copy-number ratio of the *n*-th reconstructed cycle as:

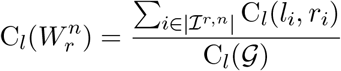

Then, rank-ordering the *N* cycles returned by a cycle decomposition algorithm in descending order, we define the *k*−heaviest copy-number ratio as the sum of the *k* largest cycles (where 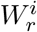 represents the *i*-th largest cycle):

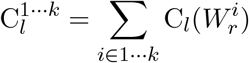

